# A universal dynamic routing architecture governs the flow of human cortical activity

**DOI:** 10.1101/2025.09.05.674413

**Authors:** Matteo Vinao-Carl, Robert Peach, Adam Gozstolai, Michael C.B David, Emma-Jane Mallas, Danielle Kurtin, Ketevan Alania, Valeria Jaramillo, Ines R. Violante, David J. Sharp, Nir Grossman

## Abstract

Behaviour requires distributed neural processing to be flexibly integrated and segregated, which in turn demands that information be dynamically routed across the brain. However, whether a unifying principle governs the routing of neural activity remains unknown. Here, we report that the flow of cortical activity is directed through canonical routing modes that are conserved across individuals, robust to variations in age, frequency band, brain state, and even the presence of neurodegeneration. These modes are constructed from the divergence and vorticity of the time-varying unit-phase vector field derived from electroencephalogram (EEG) recordings. We show that the canonical routing modes are flexibly combined to generate distinct, state- and function-dependent flow architectures that remain consistent across individuals. Disruption to the routing mode dynamics mechanistically links structural abnormalities and cognitive impairment in Alzheimer’s disease. Together, these canonical routing modes provide a unified framework for describing, modelling, and predicting the dynamic integration and segregation of macroscopic brain activity underpinning behaviour.

## Introduction

A central aim of systems neuroscience is to identify the principles by which distributed neuronal populations coordinate to support human cognition^1,2^. Since the early 2000s, network neuroscience has approached this problem by representing the brain as a graph of functionally specialized nodes that exchange information along the structural connectome^3–7^. Yet after nearly a quarter-century, it is clear that a strictly discrete network description cannot capture the full complexity and causal structure of neural dynamics^8,9^. Functional-connectivity (FC) analyses summarize pairwise statistical dependencies, leaving higher-order interactions—spanning triples and larger assemblies— unmodeled despite growing evidence from micro- and macro-scale studies that these interactions are essential for characterizing brain activity^10–12^. Standard network-based approaches are also anchored to a single spatial, temporal, and topological scale, whereas behavior depends on coordination across scales^13^. Moreover, behaviour requires the flexible redirection and routing of signals between cognitive systems^14,15^, yet conventional FC measures are undirected and therefore blind to the causal architecture governing directed information flow^16^. Together, these limitations motivate frameworks that move beyond pairwise, single-scale, and undirected descriptions of brain function.

Over the past two decades, increasingly sophisticated analytic methods have emerged to address these limitations. They include (i) higher-order-interaction tests^17–19^ and hypergraph approach to encode multi-node relations^10,20^; (ii) data-driven and model-based methods to characterise directed or “effective” communication^21–23^; and (iii) analytic techniques to explore the relationship between multi-scale network structure and behaviour ^24–29^. However, exhaustively mapping the parameter space of interaction order, direction, and scale remains prohibitively expensive, keeping an integrated study of these properties and their relationship to human behaviour underexplored.

An alternative to network-based frameworks is to model macroscopic brain activity as a vector field, with each point encoding the direction and magnitude of signal flow^30^. As in fluid systems where a continuous field emerges from countless intermolecular interactions, the extreme density and overlap of microscopic white-matter fibres in the brain yields a continuous field of neuronal activity at the macroscale^31^. Flow fields are a powerful framework for analysing neural systems because they unify three perspectives typically analysed in isolation: (i) Higher-order interactions are found around fixed points (sources, sinks, saddles, spirals) arising from nonlinear coupling among neighbouring regions^32^; (ii) directed communication can be inferred by tracing the vector streamlines ^33^; and (iii) multiscale communication can be computed through eigendecomposition of the field into patterns of signal flow spanning from low to high spatial frequencies^34^.

Over the past decade, neural flow dynamics have been linked to diverse functions, including perception, memory, and motor control^35–37^. A central phenomenon is the travelling wave (TW), which manifests as a variety of spatio-temporal patterns such as spirals^38–40^, sources and sinks^41–43^, and planar waves^44^. TWs are non-stationary their detection typically relies on estimating spatial and temporal gradients of the signals after band-pass filtering to isolate a target frequency range ^33,37,45^. Applying these approaches has revealed TWs’ roles across multiple domains: in vision, they encode stimulus position, orientation, and motion^46,47^, support sensory-motor processing of saccades^48^, and bias bistable perception^49^; in primates, they emerge in the motor cortex during movement preparation^37^ and transmit reward-related signals between frontal and parietal areas^50^. In humans, they propagate seizure fronts^51^, sweep visual areas during hallucinations induced by D-methyltryptamine DMT^52^, and dominate all stages of sleep^53– 55^, contributing to memory consolidation^56,57^. Importantly, TW directionality distinguishes cognitive processes—for example, TWs flow in opposite directions during encoding and retrieval in episodic memory tasks^44^, while the orientation of rotating blood oxygen level dependent (BOLD) waves measured with functional magnetic resonance imaging (fMRI) can classify different cognitive tasks^38^. TWs are ubiquitous across scales, observed with voltage-sensitive dyes^58–61^, local field potentials (LFPs) ^37,48,50,62,63^, electrocorticography (ECoG)^54,64– 66^,electroencephalography (EEG)^53,55,67,68^, magnetoencephalography (MEG)^69–71^, and fMRI^38,49,72,73^, and theoretical work suggests they emerge naturally from the distance-dependent and time-delay structure of the brain’s wiring diagram^33,74^ and propagate along intrinsic frequency^75^ and structural connectivity gradients^71^.

Collectively, prior work shows that modelling macroscopic brain activity as a flow field is a powerful way to capture functionally relevant dynamics. Yet most empirical studies interrogate travelling waves within a fixed temporal band (e.g., α or β-bands) or at a single spatial scale, implicitly assuming scale-specific routing. By contrast, behaviour depends on dynamic information routing that flexibly integrates and segregates neural processes across space, time, and frequency. The principles by which diverse flow patterns are orchestrated across scales therefore remain unresolved—specifically, how the brain alternates between global integration (long-range coordination of distributed regions) and local segregation (fine-scale routing supporting specialised computations). Addressing this gap requires methods that recover the higher-order, directed (causal) architecture of information flow and quantify how routing motifs are conserved, coupled, and reconfigured across scales.

Here, we report that the flow of neural activity in the human cortex is organised by a conserved family of routing modes, which are dynamically superimposed to generate distinct flow architectures that differentiate brain states and cognitive functions. We developed a computational strategy that computes the unit phase vector field (normalised spatial gradient of the instantaneous phase) from high-density EEG and extracts recurrent divergence and vorticity patterns via eigendecomposition. Across three cohorts, we show that the instantaneous direction of cortical activity can be expressed as a weighted superposition of a small family of routing modes, each operating at a characteristic spatiotemporal scale. These routing modes are conserved across individuals, frequency bands, and behavioural states, and remain intact even in neurodegenerative disease. Different cognitive tasks selectively co-activate subsets of the canonical modes, yielding distinct whole-cortex routing architectures that subserve similar functions across individuals. In Alzheimer’s disease, the modes themselves are preserved but their routing dynamics are disrupted, and these disruptions covary with regional grey-matter volume and cognitive performance. To our knowledge, this is the first demonstration that the apparent diversity of cortical flow phenomena found in EEG can be explained by a global architecture flexibly assembled from a parsimonious set of conserved routing modes, akin to how large-scale connectivity dynamics in fMRI arise from the flexible assembly of conserved network structures.

## Results

### Analysing the routing dynamics of cortical activity

We developed a computational strategy to identify recurrent routing motifs in the flow of neuronal activity from high-density brain EEG recordings. At the core of this strategy is the computation of the *unit phase vector field*, which captures the instantaneous direction of neuronal activity propagation. The neural activity at each site, *y*(*t*), is first decomposed into a set of band-limited signals using band-pass filtering (**Fig. 1a i**). These band-limited signals are then represented as nonstationary sinusoids *A*(*t*) ⋅ *sin* (ø(*t*)), where the instantaneous amplitude *A*(*t*) and phase

**Fig. 1:**
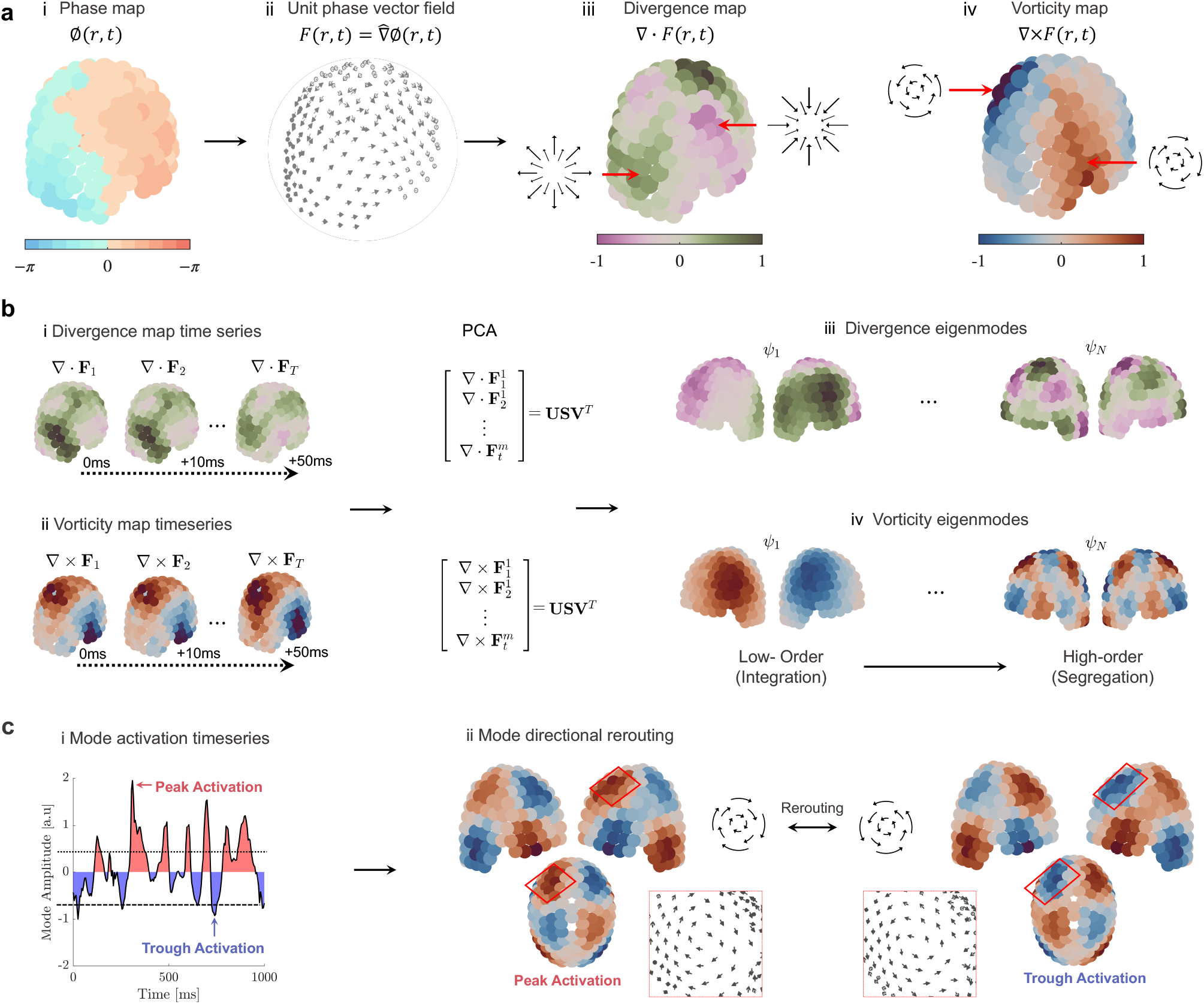
Illustration of the computation strategy. **a, Computing EEG flow routing**. *i*, Scalp EEG is band-pass filtered and the phase is computed with the Hilbert transform, yielding instantaneous phase map ∅. *ii*, Spatial derivative of the instantaneous phase map is computed approximating magnitude and direction of neural activity flow and the resulted vectors are divided by their magnitude, yielding an instantaneous unit phase vector field **F**, representing the directional routing of the flow. *iii*, Divergence and curl operators are applied to **F** to extract coherent routing changes, yielding an instantaneous divergence map (∇ · **F**) isolating source–sink flow patterns and an instantaneous vorticity map (∇ × **F**) isolating rotational flow patterns, respectively. **b, Identifying routing modes**. Principal-component analysis (PCA) is applied separately to the divergence (*i*) and vorticity (*ii*) maps, producing families of divergence eigenmodes (*iii*) and vorticity eigenmodes (*iv*), respectively. Low-order modes (*ψ*^1^) have long spatial wavelengths that support global, integrative flow, whereas higher-order modes (*ψ*^N^) exhibit finer spatial structure that partitions the cortex into segregated local flow fields. **c, Routing mode activation timeseries**. *i*, Vorticity (or divergence) maps concatenated over time are projected onto the respective eigenmodes, generating time series of mode amplitudes with peaks and troughs. *ii*, Positive and negative amplitudes correspond to topographically mirrored flow fields (e.g., clockwise rotation at a node become counter-clockwise rotational flow; source flow becomes sink flow) with each zero-crossing marks a rerouting event in cortical activity flow.

ø(*t*) are obtained via the Hilbert transform. The spatial gradient of each phase time series, ∇ø(*r, t*) is computed to yield a time-varying vector field that encodes the local direction and rate of phase progression across the cortex. The phase gradient is then normalized at each spatial location to unit length, producing the unit phase vector field 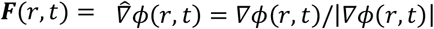, which characterizes the direction of flow independent of the strength, akin to mapping the layout of roads throughout a city while ignoring the volume of traffic (**Fig. 1a ii**). To expose coherent changes in signal routing throughout the cortex, the divergence and vorticity of the unit phase field are computed at each time point: *Div*(*r, t*) = ∇ ∙ ***F***(*r, t*), and *Vort*(*r, t*) = ∇ × ***F***(*r, t*), respectively. Divergence identifies net outflow (sources) or inflow (sinks) (**Fig. 1a iii**), while vorticity captures regions of rotational flow (clockwise or counterclockwise) (**Fig. 1a iv**) (see Arbitrary Manifold Vector Operators in Methods).

To uncover recurrent routing motifs, we concatenated the moment-by-moment divergence maps (**Fig. 1b i**) and, separately, the vorticity maps (**Fig. 1b ii**), then applied principal component analysis (PCA) to each set. This produced two families of spatial eigenmodes: divergence eigenmodes 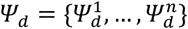 (**Fig. 1b iii**) and vorticity eigenmodes 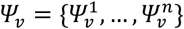 (**Fig. 1b iv**), which we collectively refer to as *routing modes*. Each routing mode represents a unique pattern of activity flow—either source/sink (divergence) or rotational (vorticity)—that recurs across time at a particular spatial frequency. The routing modes are the eigenvectors obtained from PCA of the divergence/vorticity field. Their eigenvalues form a spectrum ordered by variance explained. Projecting the maps onto these routing modes yields a time series that reflect their activation over time, with the sign of the amplitude determining the direction of flow in the underlying vector field (**Fig. 1c i**). Transitions between positive and negative amplitudes, therefore, correspond to reversals in the direction of activity flow, capturing the brain’s capacity to reroute neural signals dynamically (**Fig. 1c, ii**). This dynamic routing may be large-scale, integrative flows (low spatial frequency) to more finely segmented, localised patterns (high spatial frequency). Together, this framework represents cortical activity at each time point as a weighted superposition of routing modes, each directing the flow of activity at a particular spatiotemporal scale.

### Canonical routing modes of human cortical activity

To examine the routing dynamics of cortical activity, we recorded EEG with 127 effective electrodes in a cohort of healthy young adults (n = 19; mean age 23.0 ± 2.0 years, mean ± s.d.; 7 male, 1 non-binary, remainder female) during an overnight sleep study^76^. We analysed two resting-state sessions: one in the evening prior to sleep (session 1) and one the following morning (session 2). Each session comprised 20 minutes of EEG—10 minutes eyes open, followed by 10 minutes eyes closed—during which participants passively listened to intermittent pink noise. For further experimental details see Healthy Young Cohort in Methods.

We first computed the group-level routing modes of α-band (8–12 Hz) brain activity during the morning session by concatenating the participants’ time series of divergence and vorticity maps (separately) before applying PCA. We focused on the α-band activity given its cardinal role in the human resting state^77–79^. **Fig 2a** and **2b** show representative eigenmodes of routing vorticity and routing divergence, respectively. We found that a small subset of routing modes accounts for virtually all inter-individual variability in alpha-band routing dynamics (**Fig. 2c i**), with the first 20 vorticity modes explaining 91% of the cross-subject variance. In contrast, the first 20 divergence modes explain 96%. The evening session yields similar values of 91% and 96%, respectively (**Fig. 2f i**). We observed that vorticity routing modes were more excited at low orders compared to divergence modes (*ψ*_d1–3_, largest *P* = 4.8×10^−5^) and suppressed at high orders (*ψ*_d>15_ max *P* = 1.1 × 10^−7^) across both morning and evening sessions (**Fig 2c ii** and **Fig 2f ii**).

**Fig. 2:**
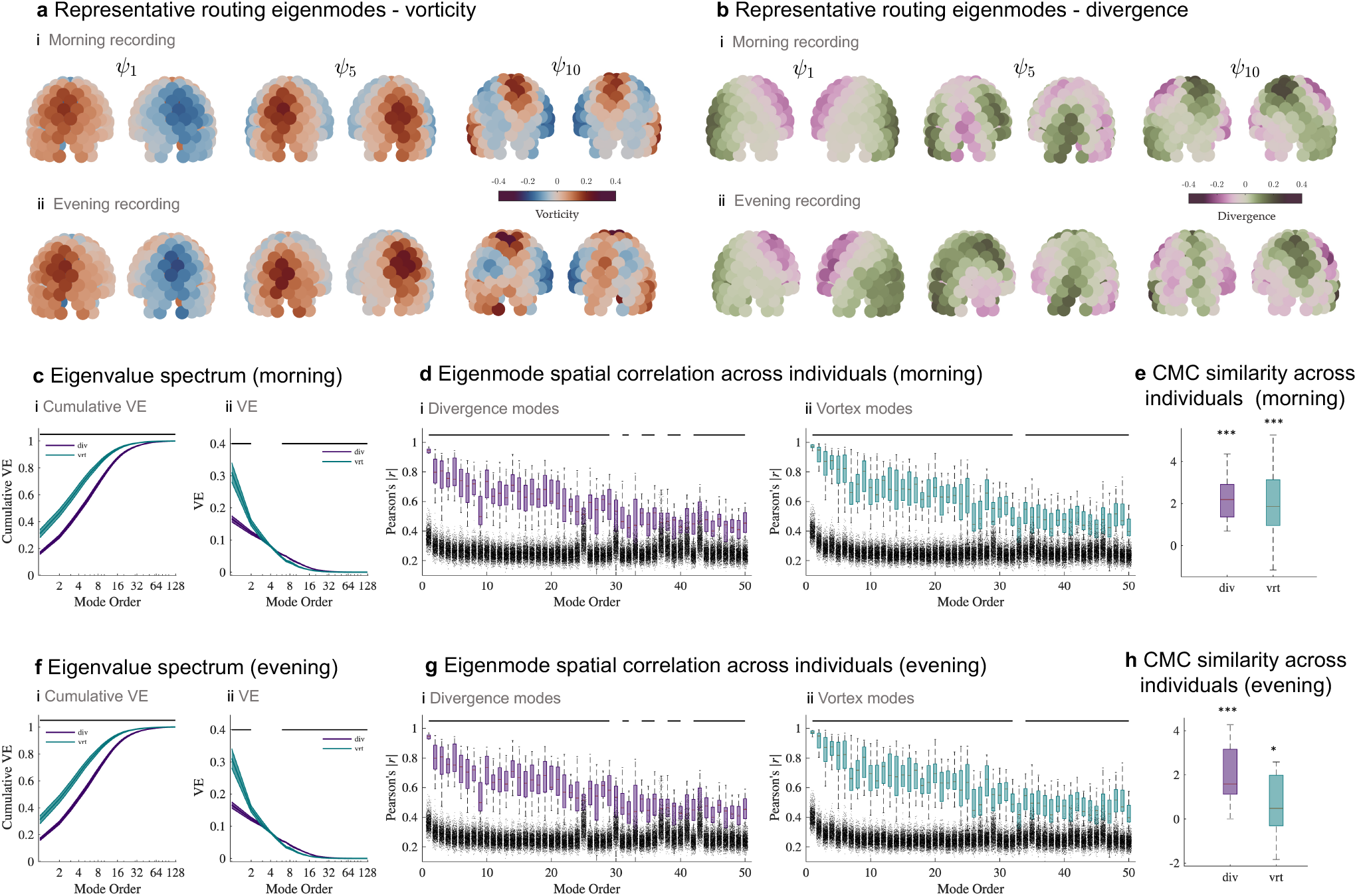
Canonical routing modes of cortical activity flow in young adults. **a–b, Representative routing eigenmodes (alpha band)**. Group-level vorticity eigenmodes (a) and divergence eigenmodes (b) from the morning EEG recording (i) and evening EEG recording (ii), showing low-order (*ψ*_1_), medium-order (*ψ*_5_), and high-order (*ψ*_10_) modes. **c, Eigenmode spectrum - morning recording**. *i*, Mean cumulative variance explained as a function of mode order for divergence (purple) and vorticity (green) modes (shading denotes 95% CI). Fewer vorticity modes (green) account for more variance across subjects than divergence modes (purple); black symbols indicate significant modes. *ii*, Raw variance contribution per mode shows vorticity modes dominate at low order (*ψ* < 4), while divergence modes dominate at higher order (*ψ* > 6). Horizontal line marks significant modes (*P* < 0.05, Bonferroni correction for 20 comparisons) **d, Eigenmode spatial correlation across individuals - morning recording**. Box plots of absolute Pearson correlations between each subject’s eigenmodes and the group reference for divergence (i) and vorticity (ii) modes. Grey boxes show a null model in which local spatiotemporal correlations are preserved but long-range structure is disrupted via spatial clustering and time-block permutation (see Methods). Boxes indicate interquartile range (IQR); central line, median; whiskers, 1.5× IQR. Horizontal line marks significant modes (*P* < 0.05, Bonferroni correction for 20 comparisons) **e, Similarity of eigenmodes coactivation across individuals - morning recording**. Similarity of cross-mode covariance matrices (CMC) across subjects for divergence modes (‘div’, purple) and vorticity modes (‘vrt’, green). CMC was derived by computing the covariance of the z-normalising amplitude time series for the first 20 divergence or vorticity modes. Similarity between CMC from different participants was assessed using cosine similarity and z-scored against a null distribution generated by 5000 column permutations (* *P* < 0.05; \*\**p* < 0.001). **f, Eigenmode spectrum - evening recording**. Showing the same as in c. **g, Eigenmode spatial correlation across individuals - evening recording**. Showing the same as in d. **h, Similarity of eigenmodes coactivation across individuals – evening recording**. Showing the same as in e.

To examine the similarity in the EEG activity routing across individuals, we computed the routing modes for each participant at the two recording sessions (evening and following morning) and quantified their similarity to the group-level morning routing modes (reference family) using pairwise absolute Pearson correlations. Each routing mode represents a spatial pattern of weights across EEG channels, and correlations were computed between these channel-weight vectors, effectively comparing the topography of each subject’s routing modes with the reference routing mode family. Since the order of the routing modes can vary between participants, we first reordered the individual routing modes to align with the order of the reference family (see Kuhn–Munkres algorithm in Methods). We found that the first 20 divergence and vorticity modes were highly correlated across participants in both the morning session (**Fig. 2d**, *ψ*_d1–d20_; *ψ*_*V*1–*V*20_, *P* < 0.001) and evening session (**Fig. 2g**, *ψ*_d1–d20_; *ψ*_*V*1–*V*20_, *P* < 0.001;) relative to a surrogate distribution (see Null Models for Mode-to-Mode Correlations in Methods), with lower order routing modes showing higher correlations. For example, in the evening recording, the absolute correlation of routing mode 1 across participants was Ψ_d1_ |*r*| = 0.86 ± 0.062 (mean ± standard deviation, SD) and Ψ_*v*1_ |*r*| = 0.97 ± 0.008 in the divergence and vorticity fields, respectively, and that of mode 20 was Ψ_d20_ |*r*| = 0.544 ± 0.041 and Ψ_*v*20_ |*r*| = 0.524 ± 0.057. Similarly, in the following morning recording, the correlations across participants was Ψ_d1_ |*r*| = 0.91 ± 0.039, Ψ_*v*1_ |*r*| = 0.97 ± 0.009, Ψ_d20_ |*r*| = 0.47 ± 0.053, and Ψ_*v*20_ |*r*| = 0.62 ± 0.072. Using the same procedure, we also tested whether the modes were reproducible within participants by computing the spatial similarity between morning and evening sessions and found similarly high correlations, (for example: Ψ_d1_ |*r*| = 0.86 ± 0.062, Ψ_*v*1_ |*r*| = 0.97 ± 0.008, Ψ_d20_ |*r*| = 0.54 ± 0.041 and Ψ_*v*20_ |*r*| = 0.52 ± 0.057 (see Supplementary Materials Fig 1).

We next tested whether the coordination between the routing modes is also conserved across individuals. We constructed a cross-mode covariance matrix (CMC) for each participant by projecting the divergence and vorticity maps onto their respective routing modes and computing the covariance of the time series of activations between all routing mode pairs. The CMC captures how strongly flow patterns are coupled between different spatial frequencies, thereby providing a compact, subject-specific fingerprint of multiscale flow organisation. We then quantified the cross-participant cosine similarity between the CMCs and z-scored these values against a null distribution, obtained by independently permuting the elements of the CMC vector 5000 times (see Supplementary Materials Fig 1 and Cross-Mode Covariance in Methods). We found that CMCs exhibit high similarity across individuals for the morning session (**Fig. 2e**, divergence modes, *t* = 7.2, *P* = 9.9 × 10−^7^; vorticity modes, *t* = 2.4, *P* = 0.02) and again for the evening session (**Fig. 2h**, divergence modes, *t* = 8.9, *P* = 5.0 × 10−^8^; vorticity modes, *t* = 5.6, *P* = 2.4 × 10−^5^), demonstrating that the multiscale flow architecture of divergence and vorticity routing is conserved across subjects.

### Canonical routing modes are conserved with age and across frequency bands

We next asked whether the routing architecture we observed in young adults persists in older age and generalises to an active task state. We recorded high-density EEG in a healthy older cohort (191 electrodes; n = 28; mean age 77.4 ± 4.9 years, mean ± s.d.; 15 women) during 5 min eyes-closed resting state and a three-session auditory attention task in which sequences of spoken letters (B, K, and M) were presented while the participants were instructed to listen passively (P task), respond to the frequent letter (F task), or respond to the oddball letter M (O task). The session order was counterbalanced between participants. Structural MRI was also acquired but not used here. See Healthy Older Cohort and Alzheimer’s Disease Cohort in Methods for experimental details.

We first examined the routing dynamics during the resting state as before but extended the analysis to three bands— low frequency (2–8 Hz, LF), mid frequency (9–20 Hz, MF), and high frequency (21–35 Hz, HF). We computed the routing modes for each participant in each band and then quantified their similarity to a reference mode family, i.e., the group-level routing modes at LF, using pairwise absolute Pearson correlations as before. **Fig. 3a** shows the reference eigenmode family and individual participant family of vorticity eigenmodes at the LF band, and **Fig. 3b** shows the corresponding spatial correlation between the individual and the reference family (reference channel weights vs subject channel weights for each routing mode). Consistent with the findings in the young cohort, the routing modes were highly correlated across individuals in all the frequency bands for the divergence eigenmodes (**Fig. 3c**, *ψ*_d1–d20_, *P* < 0.001) and vorticity eigenmodes (**Fig. 3d**, *ψ*_*V*1–*V*20_) relative to a surrogate distribution.

**Fig. 3:**
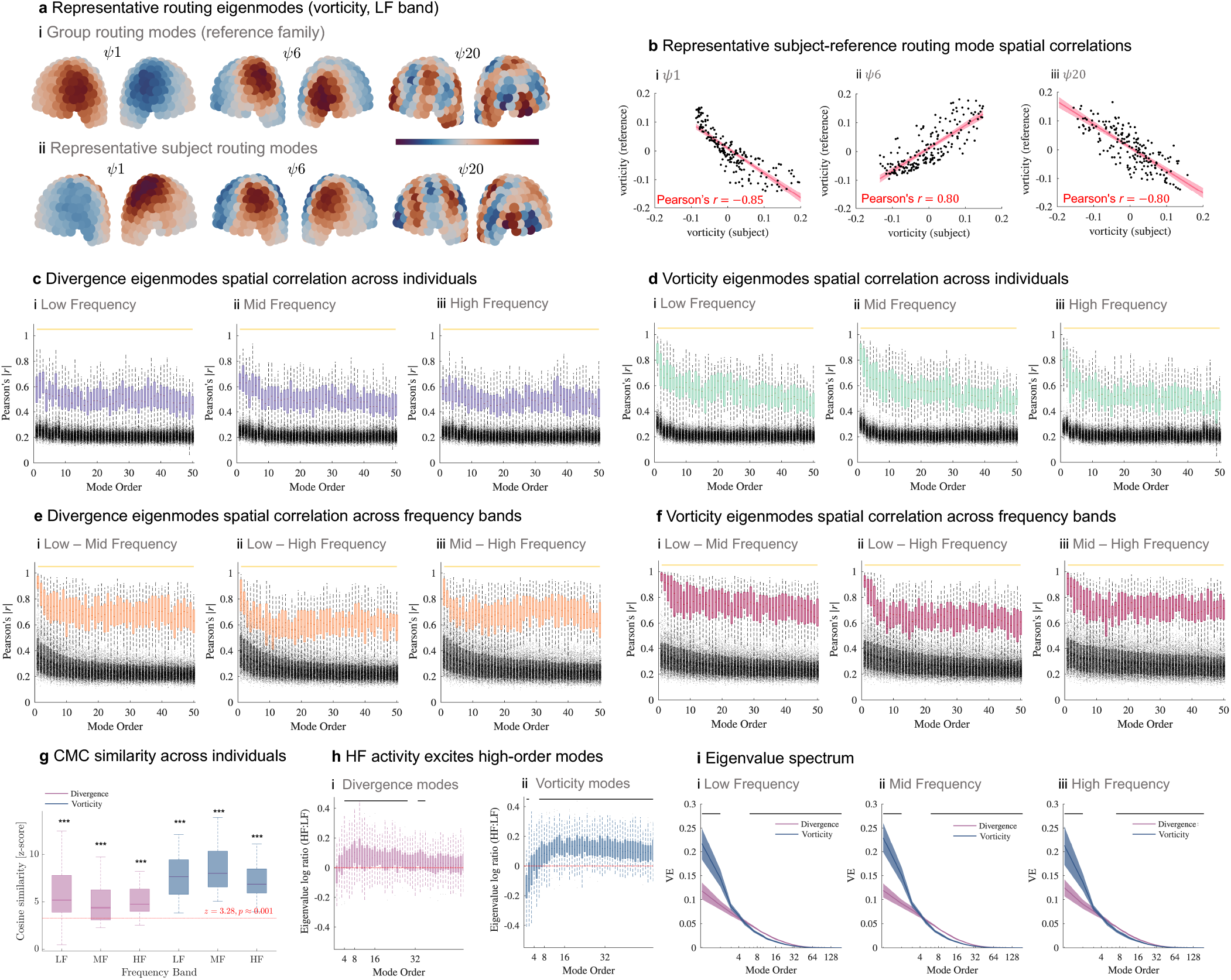
Canonical routing modes are preserved in healthy elderly individuals. **a, Representative vorticity eigenmodes (LF band)**. (i) Group-level vorticity eigenmodes (reference mode family) and (ii) Corresponding eigenmodes of a representative subject, showing low-order (*ψ*_1_), medium-order (*ψ*_6_), and high-order (*ψ*_20_) modes. CoIour scaIe bar ±1. **b, Corresponding eigenmode spatial correlations**. Group modes (reference) vs subject modes shown in (a) and the resulting spatial correlations between channel weights. Red line shows the best-fit linear regression (shaded 95 % CI). Pearson correlations: *ψ*_1_, *r* = –0.85, *P* = 1.0 × 10^−55^; *ψ*_6_, *r* = 0.80, *P* = 6.6 × 10^−44^; *ψ*_20_, *r* = –0.80, *P* = 2.9 × 10^−44^. **c, Divergence eigenmode spatial correlation across individuals**. Box plots of absolute Pearson correlations between each subject’s eigenmodes and the group reference for the first 25 modes in (i) low frequency (LF) band, (ii) mid frequency (MF) band, and (iii) high frequency (HF) band. Grey boxes show a null model in which local spatiotemporal correlations are preserved but long-range structure is disrupted via spatial clustering and time-block permutation (see Methods). Boxes indicate IQR; central line, median; whiskers, 1.5× IQR. Horizontal line marks significant modes (paired t-test, *P* < 0.05, Bonferroni correction for 20 comparisons) **d, Vorticity eigenmode spatial correlation across individuals**. Same as c but for vorticity eigenmodes. **e, Divergence eigenmodes spatial correlation across frequency bands**. Same as c but for correlations between LF vs MF modes (i), LF vs HF modes (ii) and MF vs HF modes (iii). **f, Vorticity eigenmodes spatial correlation across frequency bands**. Same as e but for vorticity eigenmodes. **g, Similarity of eigenmodes coactivation across individuals**. Similarity of CMC across subjects for divergence modes (purple) and vorticity modes (blue) in the LF, MF and HF bands for divergence (purple) and vorticity (blue). Red dashed line, paired t-test; Z = 3.28, *P* = 0.001) **h, Eigenvalue frequency band ratio**. Log ratio of the eigenvalue in the HF band and the LF band for the first 50 routing modes (mean ± 95 % CI). (i) Divergence modes, (ii) Vorticity modes. Negative values indicate stronger expression in the LF band; positive values indicate stronger expression in the HF band. Horizontal line marks significant modes (paired t-test, *P* < 0.05, Bonferroni correction for 50 comparisons). **i, Eigenvalue spectrum**. Mean cumulative variance explained as a function of mode order for divergence (purple) and vorticity (blue) modes (shading denotes 95% CI). (i) LF band, (ii) MF band, and (iii) HF band. Vorticity dominates low-order modes (*ψ*_1–2_), while divergence contributions are greater in higher-order modes (*ψ*>8). Black markers indicate statistically significant differences (paired t-test, *P* < 0.05, Bonferroni correction for 127 comparisons).

Within individuals, routing modes were conserved across frequency bands. Spatial patterns of both divergence (**Fig. 3e**, *ψ*_d1:50_) and vorticity modes (**Fig. 3f**, *ψ*_*v*1:50_) were highly correlated between LF, MF, and HF bands. For example, vorticity mode 1 showed strong correlations across bands (LF v MF *ψ*_*v*1_ |r| = 0.95 ± 0.02, LF v HF *ψ*_*v*1_ |r| = 0.90 ± 0.037;; MF v HF *ψ*_*v*1_ |r| = 0.90 ± 0.038), while higher-order routing modes such as mode 25 remained significantly correlated, albeit at lower magnitudes (LF–MF: *ψ*_*v*25_ |r| = 0.73 ± 0.050; LF-HF *ψ*_*v*25_ |r| = 0.64 ± 0.054; MF–HF: *ψ*_*v*25_ |r| = 0.72 ± 0.039, all *P* < 10^−5^, Bonferroni-corrected). These findings indicate that a single family of routing modes recurs across frequency bands.

As in the young cohort, we tested whether multiscale coordination between routing modes is reproducible across individuals. For each frequency band, we quantified multiscale coordination by computing the CMC matrix, which captures the temporal covariance across modes of different orders. We then compared the similarity between participants’ CMCs for divergence modes (purple) and vorticity modes (green). CMC similarity was significantly greater than expected by chance across all bands: divergence modes, LF t = 10.22, *P* = 8.9 × 10^−11^; MF t = 12.17, *P* = 1.8 × 10^−12^; HF t = 16.10, *P* = 2.3 × 10^−15^; vorticity modes, LF t = 18.41, *P* = 8.2 × 10^−17^; MF t = 18.09, *P* = 1.3 × 10^−16^; HF t = 17.52, *P* = 2.8 × 10^−16^ (**Fig. 3g**). These results indicate that the routing architecture governing cortical activity is conserved in older adults and invariant across frequency bands.

Oscillatory activity is known to exhibit a well-established frequency–scale relationship: slow rhythms recruit widespread cortical territories, whereas high-frequency activity is spatially focal^80–83^, exhibiting distinct spatial patterns ^84^. This appear at odds with our finding of a single routing mode family governing activity flow across all frequency bands. We hypothesized that shifts in the relative weighting of low-order (global, integrative) versus high-order (local, segregative) modes could bias the cortex toward global or local routing regimes. To test this we matched routing modes across low and high bands by spatial similarity using the Kuhn–Munkres algorithm. For each matched pair, we computed the eigenvalue ratio (HF:LF) to index the mode’s relative weighting at high versus low frequencies. We found that the routing mode eigenvalue ratio increased with frequency: low frequencies preferentially engaged low-order, global routing modes, whereas high frequencies recruited higher-order, local routing modes (**Fig. 3h**). As in young adults, vorticity modes predominated at low orders and divergence modes at higher orders across all bands (**Fig. 3i**). These results delineate two complementary control axes for tuning integration versus segregation: (i) the relative weighting between vorticity and divergence eigenmodes and (ii) the relative weighting of low-versus high-order modes within each frequency band.

### Cognitive tasks evoke consistent routing architectures across subjects

We next examined the routing of neural activity during an auditory attention task (**Fig. 4)**. We compared the routing modes of individual participants at the LF, MF, and HF bands across the entire task (by separately concatenating the sessions’ time series of divergence and vorticity maps before performing PCA) and quantified the similarity to the same reference routing mode family as before (group-level resting-state LF routing modes). We found that the routing modes during the task were highly correlated to those during rest, with a higher absolute correlation at lower order routing modes, **Fig. 4a** (for example: 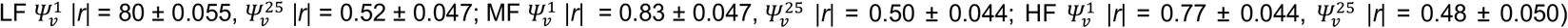, confirming that the canonical routing architecture persists across brain states (for divergence eigenmodes see Supplementary Materials Fig 2). However, task engagement shifted the routing architecture towards stronger activation of high-order modes, particularly at the high-frequency band (**Fig. 4b**).

**Fig. 4:**
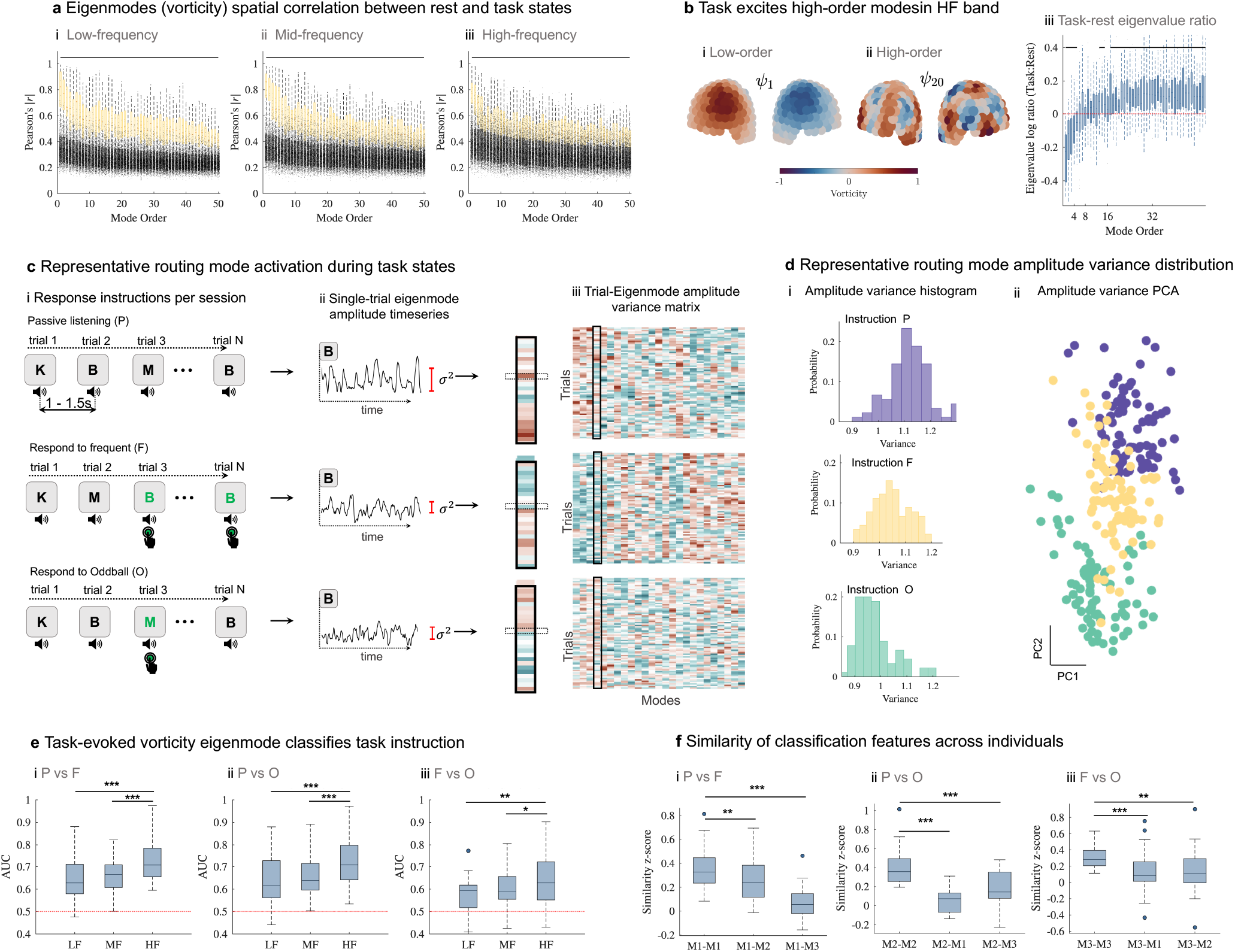
Cognitive tasks evoke consistent routing architectures across subjects. **a, Vorticity eigenmode spatial correlation between rest and task states**. Box plots show absolute Pearson correlations between rest and task eigenmodes within each participant for (i) low-frequency (LF), (ii) mid-frequency (MF), and (iii) high-frequency (HF) bands. Grey boxes show surrogate null distributions in which local per-channel statistics were preserved but cross-channel temporal dependencies were disrupted by channel-wise substitution (see Null Model for Mode-to-Mode Correlation in Methods). Boxes indicate interquartile range (IQR); central line, median; whiskers, 1.5× IQR. Horizontal line marks significance threshold, defined as the 95th percentile of the surrogate null distribution (P < 0.05, Bonferroni corrected for 50 comparisons). **b Eigenvalue task-rest ratio**. (i) Representative low-order mode (ii). Representative high-order mode. (iii) Boxplots show the group-level distribution of the log_10_ eigenvalue ratio (task vs rest) for modes *ψ*_1-50_. Error bars indicate 95% confidence intervals; black markers denote modes with significant change relative to rest (paired t-test, *P* < 0.05, uncorrected). **c, Representative task-evoked routing mode activation**. (i) Auditory attention task with three sessions that differ by instruction: passive listening (“P”), respond to frequent stimulus (“F”) and respond to oddball stimulus (“O”). (ii) Eigenmode amplitude timeseries evoked by the letter ‘B’ during passive frequent and oddball task instructions. Red bars indicate mode amplitude variance. (iii) Trials × eigenmodes activation amplitude matrices during the tasks: “P” (Top matrix), “F” (middle matrix) and “O” (bottom matrix). **d, Representative routing mode amplitude variance distribution**. (i) Histogram of eigenmode activation amplitude across trials during the tasks shown in c. (ii) PCA of the trial-level eigenmode activation in c, showing clustering of amplitude variance by “P”, “F” and “O” instructions despite identical stimuli. **e, Task-evoked vorticity eigenmode classifies task instruction**. Binary classification of task instruction on single trials using the vorticity eigenmode amplitude variance as a predictor, showing area under the curve (AUC) ± 95% CI across participants separately for LF, MF, and HF bands. HF modes yield the highest classification AUC: P vs F, 0.75 ± 0.04 (i); P vs O, 0.74 ± 0.04 (ii); F vs O, 0.68 ± 0.04 (iii). *P* < 0.05, *< 0*.*01*, < 0.001 (paired t-test, uncorrected). **f, Similarity classification features across individuals**. Cosine similarity of the classifier feature-importance vector across participants, showing that matched comparisons (same task contrast, different individuals) are significantly higher than mismatched comparisons, for P vs F (i), P vs O (ii), and F vs O (iii), indicating that similar subsets of routing modes encode task distinctions across subjects *P* < 0.05, < 0.01, < 0.001 (paired t-test, uncorrected).

To test whether different behaviours activate district routing architectures, we examined single-trial mode activations under the different auditory attention rules (i.e., passive listening (P), respond to frequent (F) stimuli, and respond to oddball (O) stimuli) (**Fig.4c i)**. Divergence and vorticity maps for each participant were projected onto the first 25 group-level routing modes (**Fig.4c ii)**, and amplitude variance for each routing mode was computed in the 1-second window following stimulus onset, yielding a trial-by-mode amplitude variance matrix (**Fig.4c iii**). We observed that the mode amplitude variance distribution differed depending on the attentional rule, as is evident in the representative mode-amplitude distribution for each task instruction in **Fig 4d**.

To test whether different behaviours during the task elicited distinct routing architectures we trained a binary random forest classifier on the 25 mode-amplitude values per trial (see Classification of task session using random forests in Methods). We found that despite the identical auditory input, the task rule was accurately classified from the mode amplitude variance on single-trials (**Fig. 4e**). The highest AUC was achieved when using the HF vorticity eigenmodes (Passive vs Frequent = 73.3% ± 3.9%; Passive vs Oddball = 73.0% ± 3.8%; Frequent vs Oddball = 68.0% ± 4.5%) and for HF divergence eigenmodes (Passive vs Frequent = 64.4% ± 4.0%; Passive vs Oddball = 74.2% ± 4.1%; Frequent vs Oddball = 68.0% ± 3.8%, see Supplementary Materials Fig 2c). Participant performance was at ceiling across all three tasks (Passive: 97.5% ± 12.5%; Frequent: 98.2% ± 3.3%; Oddball: 97.5% ± 6.1%). Pairwise comparisons confirmed no significant differences (PvO: *t* = –0.006, *P* = 0.99; FvO: *t* = 0.51, *P* = 0.61; PvF: *t* = –0.28, *P* = 0.09), ruling out performance differences as a confounding factor.

We next asked whether the routing architectures evoked by the attentional tasks were consistent across individuals. To test this, we extracted feature-importance vectors from each participant’s three Random Forest classifiers (Passive vs Frequent, Passive vs Oddball, Frequent vs Oddball). Each classifier assigned an importance weight to every routing mode, yielding a vector that summarised which routing modes carry the most information for distinguishing between task instructions in that individual. To assess whether participants rely on the same subset of routing modes when attending to the same instruction, we compared feature-importance vectors across individuals in two ways: *matched pairs*, where the same model was compared between two participants (e.g., Alice–Model 1 vs Bob–Model 1), and *mismatched pairs*, where different models were compared across participants (e.g., Alice–Model 1 vs Bob–Model 2). For each pairwise comparison, we computed the absolute cosine similarity between the corresponding feature-importance vectors: values range from 0 to 1, with values closer to 1 indicating a more substantial alignment. We found that, across all three models, matched comparisons consistently showed higher cosine similarity than mismatched ones (**Fig. 4f**). For vorticity eigenmodes, matched pairs in Model 1 exceeded Model 1 vs Model 2 (t = 3.2, p = 0.0033) and Model 1 vs Model 3 (*t* = 7.9, *P* = 1.6 × 10^−8^); matched pairs in Model 2 exceeded Model 2 vs Model 1 (*t* = 11.3, *P* = 1.0 × 10^−11^) and Model 2 vs Model 3 (*t* = 6.8, *P* = 2.8 × 10^−7^); and matched pairs in Model 3 exceeded Model 3 vs Model 1 (*t* = 4.6, *P* = 8.7 × 10^−5^) and Model 3 vs Model 2 (*t* = 3.7, *P* = 9.8 × 10^−4^), for results with divergence modes, see Supplementary Materials Fig 2). In other words, when two individuals attend to the same instruction, the flow of their cortical activity is routed more similarly than if they were attended to different rules.

### Neurodegenerative pathology leaves routing modes intact

Having established that the flow of cortical activity is moulded by a set of canonical routing modes that are conserved in healthy individuals, we next examined the routing dynamics under a disease condition in which the neural tissue is compromised due to neurodegeneration. We repeated the experiment in a cohort of participants with moderate Alzheimer’s disease (AD; n = 33, 18 females; age = 76.3 ± 7.8 years), matched in age to the healthy older cohort (no significant age difference, *P* = 0.51). The Alzheimer’s Disease Assessment Scale - Cognitive subscale (ADAS-Cog)^85^ was 37.1 ± 12.1, suggesting moderate impairment. The patients showed reduced accuracy in the simple attention tasks relative to the healthy cohort (93.3 ± 7.6% vs. 97.8 ± 3.3%, *t* = –2.9, *P* = 0.0052, Supplementary Materials Fig 3).

We applied the computational framework to the band-limited EEG to extract subject divergence and vorticity eigenmodes in AD and tested whether the modes were preserved relative to the healthy older cohort by computing the similarity of each participant’s routing modes to the reference mode family (group-level resting-state LF routing modes in the healthy older cohort). During the resting state, we found that the low and high order routing modes remained remarkably intact in the AD cohort (**Fig. 5a,b**). Resting modes across all frequency bands were highly correlated with the reference routing mode family in the healthy older cohort for both divergence eigenmodes **(Fig. 5c)** and vorticity eigenmodes (**Fig. 5d)**, and these routing modes were also conserved between rest and task states (Supplementary Materials Fig 4).

**Fig. 5:**
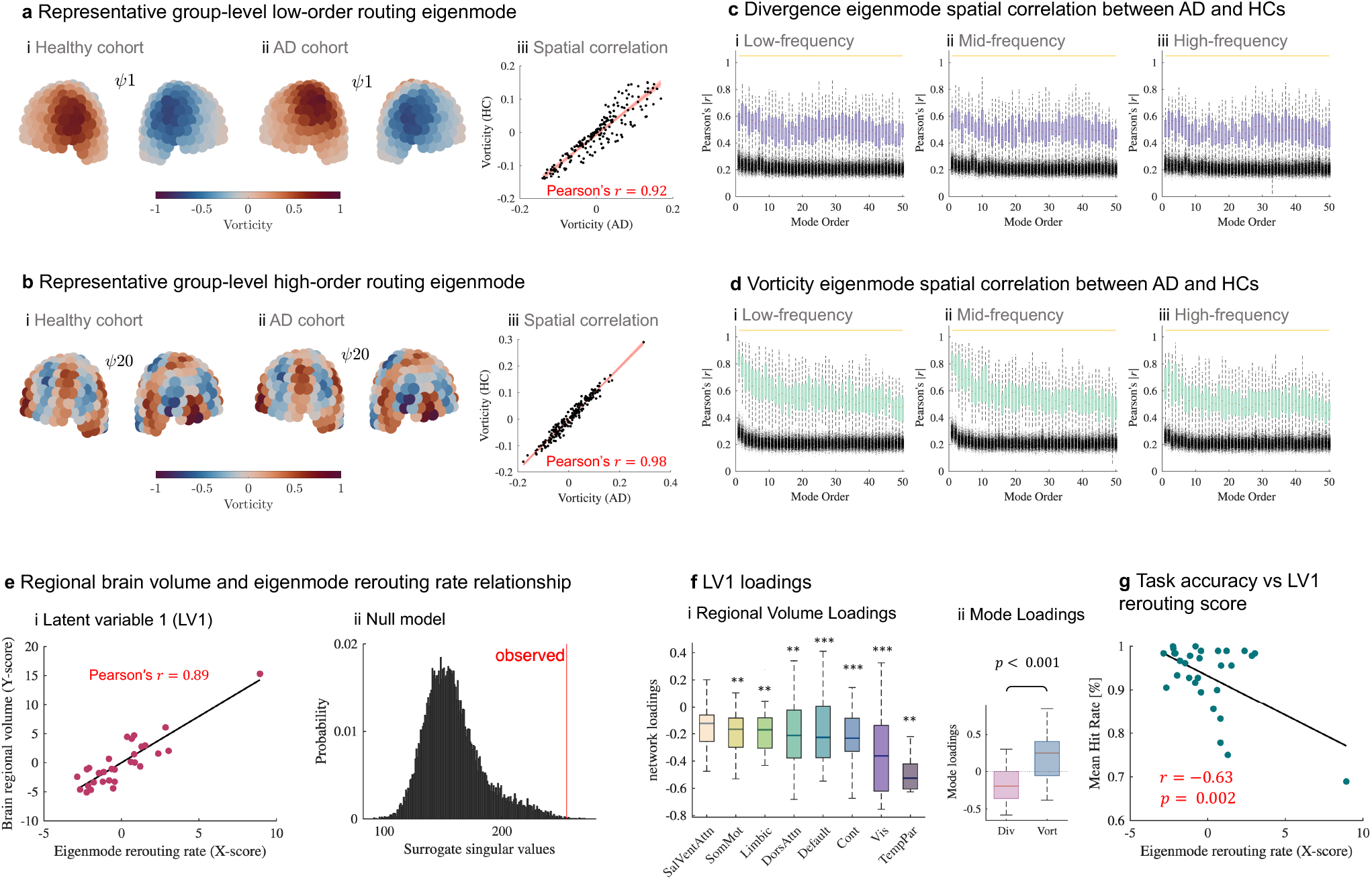
Cortical activity routing in Alzheimer’s disease. **a, Representative group-level low-order routing eigenmode (LF band, rest)**. (i) Vorticity eigenmode *ψ*_1_ in the healthy older cohort (HC), (ii) the same eigenmode *ψ*_1_ in the AD cohort, (iii) Corresponding eigenmode spatial correlations between the HC and AD cohorts in i and ii. Red line shows the best-fit regression (shaded 95 % CI). Pearson’s *r* = 0.92. **b, Representative group-level high-order routing eigenmode (LF band, rest)**. Showing the same as a but for *ψ*_20_. Pearson’s *r* = 0.98. **c, Divergence eigenmode spatial correlation between AD and healthy cohorts**. Box plots of absolute Pearson correlations between HC reference family eigenmodes and AD routing modes in (i) LF band, (ii) MF band, and (iii) HF band. Grey boxes show a null model in which local channel structure is preserved but long-range correlations disrupted randomly sampling vorticity /divergence time series from permutation (see Null Model for Mode-to-Mode Correlation in Methods). Boxes indicate IQR; central line, median; whiskers, 1.5× IQR. Horizontal line marks significant modes (*P* < 0.05, Bonferroni correction for 50 comparisons). **d, Vorticity spatial correlation between AD and healthy cohorts**. Same as c but for vorticity eigenmodes. **e, Regional brain volume and eigenmode rerouting-rate relationship**. Partial least squares (PLS) analysis for correlation between rerouting rate matrix (X) and brain volume matrix (Y). (i) In-sample correlation between latent variable 1 (LV1) rerouting-rate scores and regional brain volume scores, showing a strong shared co-variance across participants (Pearson’s *r* = 0.89), accounting for 20 % of the total cross-covariance between X and Y. **f, LV1 loadings**. (i) LV1 loadings for regional brain volume, group by canonical networks — Salience/Ventral Attention (SalVentAttn), Somatomotor (SomMot), Limbic, Dorsal Attention (DorsAttn), Default Mode (Default), Frontoparietal Control (Cont), Visual (Vis), and Temporo-parietal (TempPar) — and (ii) loadings for divergence (purple) and vorticity (blue) modes. Larger grey-matter volume is associated with faster re-routing of vorticity modes and slower re-routing of divergence modes (paired t-test, *P* <0.001). Boxes indicate IQR; central line, median; whiskers, 1.5× IQR **g, Task accuracy vs LV1 rerouting score**. Pearson’s correlation between the patients’ LV1 rerouting rate scores (X-scores) and their mean performance accuracy across oddball and frequent sessions in the attention task (Pearson’s *r* = –0.63, *P* = 0.002).

Prior studies have suggested that at this stage of disease progression, cortical activity becomes less flexible, with slower temporal dynamics at low frequencies ^86–89^ and reduced entropy and complexity ^87,90,91^ This points to a potential mechanistic shift: structural pathology may constrain the flexibility with which cortical activity is rerouted. To test this, we quantified mode rerouting rates in the LF band—defined as the frequency of amplitude zero-crossings per second during rest—as a proxy for the dynamic adaptability of cortical flow routing at low frequencies. For each patient, we computed the rerouting rate of the first 25 LF vorticity and 25 LF divergence eigenmodes during resting state and extracted the volume in 214 cortical and subcortical regions (see Partial Least Squares Analysis in Methods), resulting in a 30 × 50 activity rerouting rate matrix (X) and 30 × 214 brain volume matrix (Y). We then deployed a partial least squares (PLS) analysis to identify patterns of covariance between the rerouting rate and brain volume matrices.

PLS revealed a single significant latent variable pair (LV1), comprising an X score (linear combination of LF routing modes) and a Y score (linear combination of regional grey-matter volumes) that were strongly correlated (*r* = 0.89, *P* < 0.0001, **Fig. 5e**). Grey-matter volume was significantly associated with rerouting rate in Salience/Ventral Attention (SalVentAttn: *t* = −3.17, *P* = 0.0073), Somatomotor (SomMot: *t* = −3.93, *P* = 0.0073), Limbic (Limbic: *t* = −4.08, *P* = 0.0008), Dorsal Attention (DorsAttn: *t* = −4.28, *P* = 0.0002), Default Mode (Default: *t* = −4.65, *P* = 0.0002), Control (Cont: *t* = −6.50, *P* = 0.0001), Visual (Vis: *t* = −6.52, *P* < 10^−4^), and Temporo-parietal networks (TempPar: *t* = −7.79, *P* = 0.0006; **Fig. 5f, i**). The loadings of LV1 were consistently positive for vorticity eigenmodes and negative for divergence eigenmodes (**Fig. 5f, ii**), indicating that larger grey-matter volumes correspond to faster rerouting rates of vorticity eigenmodes and slower rerouting of divergence eigenmodes (paired t-test, *P* < 0.001). Importantly, the patients’ LV1 rerouting rate scores (X-scores) were negatively correlated with their performance in the attention tasks (Pearson’s *r* = –0.62, *P* = 0.002; **Fig. 5g**). Together, these findings indicate that cognition in early stages of AD may falter not because spatial communication pathways are lost, but because the capacity to reroute the flow of activity along preserved pathways becomes disrupted.

## Discussion

This paper presents a computational strategy to characterise the routing of neural activity across multiple spatio-temporal scales of the human cortex. We show that cortical activity is routed through a set of canonical modes that are dynamically superimposed to create distinct flow architectures depending on brain state and cognitive function. We used the gradient of the instantaneous phase to approximate spatial change (i.e., flow) of neural activity consistent with earlier studies^33,37,71,73^. However, here we normalised the resulting phase vectors by dividing by the magnitudes. The resulting unit phase vector field allows for isolating the backbone of neuronal signal flow—the brain’s routing geometry, independent of transient fluctuations in the strength, akin to mapping the layout of roads throughout a city while ignoring the volume of traffic. Future work may explore the relationship between the routing architecture derived from the unit phase vector field and the activity flow itself, extracted from the unnormalized phase vector field. We next used divergence and curl operators to expose coherent changes in signal routing. The resulting divergence map captures pathways of outflow (source) or inflow (sink), while the vorticity map captures rotational flow (clockwise or counterclockwise). Regions with low divergence and vorticity have minimal change in the routing direction. The separation between divergence and vorticity routing changes is artificial. In fact, several divergence-vorticity eigenmode pairs have highly correlated time courses and comparable predictive power for both task classification in healthy subjects and behavioural outcomes in Alzheimer’s disease, suggesting partial redundancy and an effective dimensionality of the cortex lower than the 50 vorticity and divergence routing modes reported here. Joint dimensionality-reduction across both families could clarify this further. We used PCA to uncover recurrent routing motifs over time in the time series of divergence and vorticity maps (instantaneous maps were concatenated). The use of PCA has proven efficient in uncovering consistent patterns across individuals and datasets, but it is not unique. It is plausible that other dimensionality reduction approaches, such as independent component analysis^92^, dynamic mode decomposition^93^ or empirical mode decomposition^94^ may also work well.

A central aim of systems neuroscience is to identify latent representations that capture how the brain coordinates neural activity to support cognition. The neural network framework, described by a parsimonious set of functionally linked regions, proposes that neural information is routed through selective pathways between the networks. Yet today, it is increasingly clear that the discrete network framework cannot capture the full complexity, higher-order structure, and causal dynamics of neural activity. The contemporary neural flow field framework overcomes these limitations. Our findings suggest that the direction of neural flow dynamics across the cortex can be reconstructed from a parsimonious set of activity routing eigenmodes, as theoretically predicted by neural field models ^95,96^albeit not routing eigenmodes. Earlier studies have demonstrated that spatial characteristics of brain activity can be reconstructed from eigenmodes derived from the structural characteristics of the brain, such as the connectome ^97,98^ or geometry^95^. Pang et al. derived geometric eigenmodes of the brain by applying the Laplace–Beltrami operator to cortical surfaces reconstructed from structural MRI and demonstrated that these geometric eigenmodes can approximate the spatial distribution of functional connectivity during rest and average task-evoked activity in the HCP dataset^99^. Overall, canonical structural eigenmodes have been shown to successfully capture spatial maps of brain activity, but they do not represent its causal spatiotemporal flow. In contrast, the canonical routing eigenmodes introduced here are distinct, as they are derived directly from directional EEG time series and thus intrinsically capture the directed spatiotemporal dynamics of brain activity.

A potential objection is that any eigendecomposition yields a complete set of orthogonal basis functions capable of reconstructing any smooth signal, so the patterns we extract might be purely mathematical artefacts. Several observations rebut this. First, we find that mode topography is nearly identical across participants, and similar modes are consistently activated during the same behaviours. On an arbitrary basis, there is no reason to expect two individuals to share the same set of routing modes, let alone recruit them for the same cognitive operations. Robust global structures have been reported in fMRI data where they are consistent across subjects^4,100,101^ and conserved across species^102^. Reproducible spatial patterns of activity have also been observed in the EEG as microstates^103,104^ and these components remain preserved even in neurological disease^105^. However, microstate reproducibility is usually limited to <7 components^106,107^. In contrast, we observed reproducibility of the flow-field routing modes up to high orders (*ψ* > 50), and of their functional recruitment for the first 25 modes (*ψ*1–25). Second, our decomposition consistently yields two routing mode families with distinct power spectra: vorticity modes dominate low spatial frequencies (supporting course-grained global integration), while divergence modes dominate higher spatial frequencies (supporting fine-grained local segregation). If the basis were generic, there would be no intrinsic reason for one family to systematically carry more or less power. This apparent functional dissociation suggests that the routing modes reflect fundamental organisational principles of the cortex. Third, the observation that task engagement and higher EEG frequencies preferentially up-regulate higher-order divergence modes—and suppress them at rest and in lower frequencies—is inconsistent with the behaviour of an arbitrary set of basis functions.

Neural-mass models have shown that neural flow patterns such as vortices, sources, and sinks reconfigure at brain state transitions^33^. More recently, intracranial data have shown that the direction of travelling waves during memory tasks changes depending on the memory operations^44^. Our results generalise and empirically ground these ideas in three key ways. First, our findings demonstrate that wave patterns, such as vortices, sources, and sinks, are local expressions of a global routing architecture rather than isolated phenomena. Second, we show that the modulation of activity flow direction occurs not only between encoding and retrieval, as reported in^44^, but also as subjects alternate between different task instructions, and between resting and task states. This suggests that rerouting of neural activity flow is a general control principle to alternate between brain states and cognitive functions. Third, we show that structural abnormalities and cognitive deficits are associated with changes in the rate of rerouting neural activity flow in AD, suggesting that rerouting disruption could be a mechanistic pathway through which neural atrophy leads to cognitive decline. Our findings are consistent with recent fMRI work showing that individuals whose structural and functional connectomes are more closely aligned tend to score higher on measures of general cognition^108^. Here, we find the same principle in EEG: LF modes that are highly correlated with subcortical grey matter volume also modulate attentional performance.

Classically low and high-frequency neural activity are thought to reflect global and local processing respectively ^80^, arising from two distinct mechanisms of long-range cortical interactions vs local excitatory–inhibitory interactions ^109^. Our results challenge this dichotomy. We show that neural activity at all frequencies is governed by the same canonical family of routing modes and that high-frequency neural activity can also form stable global flow patterns. These findings are consistent with recent intracranial recordings demonstrating that very high-frequency neural activity (100-400Hz) exhibits long-range spatial synchronisation patterns^110^. We extend this principle to demonstrate that every frequency band has access to both globally integrative and locally segregative flow patterns, depending on the order of the modes engaged. The key distinction is that high-frequency activity more frequently recruits higher-order (segregative) routing modes, whereas low-frequency activity is dominated by lower-order (integrative) modes. Nevertheless, the underlying canonical routing architecture constraining activity remains the same across all temporal scales.

Our results demonstrate that dynamic activation of conserved routing modes provides a principled taxonomy of how the spatial flow of activity is flexibly integrated and segregated to shape information processing in the brain. This suggests that the routing modes may represent state variables for the human cortex.

## Methods

### Healthy Young Cohort

#### Participants

18 healthy adults (7 men, 1 non-binary; 23.0 ± 2.0 years, range 18–30) completed the protocol. Volunteers were free of neurological, psychiatric or sleep disorders, maintained a body-mass-index < 30, reported no trans-meridian travel or shift work in the previous month, and were not taking medication affecting the central nervous system. For seven days before the laboratory visit, participants kept a fixed 8-h sleep schedule verified by wrist actigraphy (Actiwatch Spectrum Plus, Philips Respironics) and daily sleep diaries. Caffeine and alcohol were prohibited for 24 h prior to arrival. Recordings were performed in a sound-attenuated, temperature-controlled, windowless bedroom at the Surrey Sleep Research Centre. Participants arrived ~3 h before habitual bedtime and remained in the laboratory until the end of the 10-h sleep opportunity the following morning. Only the wake EEG blocks described below are analysed here. Written informed consent was obtained in accordance with the Declaration of Helsinki; the study was approved by the University of Surrey Ethics Committee.

#### EEG acquisition

High-density EEG was recorded with a 128-channel actiCAP-snap system (Brain Products, Germany) positioned so that Cz lay at the cap centre. Additional bilateral canthus EOG, chin EMG, and ECG electrodes were applied according to AASM guidelines. Signals were sampled at 500 Hz with 24-bit resolution and referenced to Cz during acquisition; re-referencing was performed offline as described in the Analysis section. Electrode impedances were maintained below 50 kΩ.

#### Morning and evening resting-state EEG recordings

We analysed continuous wake EEG from two resting sessions: one recorded in the evening before sleep, and one the following morning after overnight sleep monitoring. Each session consisted of two consecutive blocks of 10-minutes eyes-open EEG recording (EO) followed by 10-minutes eyes-closed (EC) during which participants remained awake. In the EO block, participants fixated on a central cross, while in the EC block they were instructed to close their eyes. Throughout both conditions, sparse auditory stimulation was presented in the form of 20-ms pink-noise bursts (1/f) at 50, 55, and 60 dB delivered binaurally via in-ear headphones with a jittered inter-stimulus interval of 1.5 ± 0.3 s (range: 1–2 s). For full methodological description see source paper^76^.

#### EEG preprocessing for all datasets

All EEG recordings were preprocessed in EEGLAB 109 (v2025.0.0) running under MATLAB R2024a. EEG signals were band-pass filtered between 2 and 40 Hz using a zero-phase, second-order Butterworth filter and subsequently downsampled to 200 Hz. Channel rejection was performed using EEGLAB’s `pop_rejchan` function, removing channels with kurtosis in the 2–35 Hz frequency range exceeding ±4 standard deviations. The remaining data were re-referenced to the common average. Independent component analysis was performed using `pop_runica` with the Picard algorithm. Components were automatically classified using the ICLabel plugin, and up to three components identified as non-neural were removed per participant.

### Healthy Older Cohort and Alzheimer’s Disease Cohort

#### Participants

A total of 42 participants with a clinical diagnosis of Alzheimer’s disease (AD) were recruited from two studies. The first was Minder, an ongoing longitudinal, community-based cohort study of individuals living with dementia, conducted by the Care, Research and Technology Centre of the UK Dementia Research Institute. Although Minder includes individuals with various dementia subtypes, only those with a diagnosis of AD were included in the present study. The second source was the Physiological Correlates of Noradrenergic Add-on Therapy (PCNorAD) study, which investigates noradrenergic dysfunction in AD. Participants in PCNorAD had previously completed the NorAD clinical trial. With the exception of one individual, all participants from NorAD were enrolled in PCNorAD no earlier than one month following trial completion, to ensure adequate medication washout before participation in this study. In addition, 32 age-matched healthy older control participants (HCs) were recruited as part of PCNorAD. All AD participants had a pre-existing clinical diagnosis of probable Alzheimer’s disease, which was re-evaluated by a multidisciplinary team (MDT) comprising neurologists, psychiatrists, and neuroradiologists. This diagnostic review integrated neuroimaging and cognitive assessment data acquired as part of the present study, alongside clinical history and prior medical investigations. Exclusion criteria for both groups included any major neurological or psychiatric condition other than AD. One HC was excluded due to evidence of traumatic diffuse axonal injury on MRI, and another was excluded due to objective cognitive impairment, defined as a score greater than 20 on the Alzheimer’s Disease Assessment Scale–Cognitive Subscale (ADAS-Cog). Two AD participants were excluded after MDT review concluded that AD was unlikely to be the primary cause of their clinical presentation. In addition, EEG data from two healthy controls and seven AD participants were excluded due to poor data quality. This yielded a final sample of 33 participants with AD and 28 healthy controls included in the EEG analyses.

#### MRI Acquisition

Structural MRI data were acquired for the majority of participants; however, six individuals did not undergo MRI scanning. Four healthy control participants declined the MRI procedure, while two patients with Alzheimer’s disease (AD) were excluded due to severe cognitive impairment and claustrophobia. The total number of AD participants with complete strucrural MRIs was 30. The maximum interval between MRI acquisition and EEG recording was one year, with a mean delay of three months.

MRI scans were performed using a Siemens Verio 3.0-T scanner equipped with a 32-channel head coil. Structural imaging was obtained using a T1-weighted three-dimensional magnetization-prepared rapid acquisition gradient-echo (3D-MPRAGE) sequence, with a total acquisition time of 5 minutes and 3 seconds. The imaging parameters were as follows: flip angle = 9°, echo time (TE) = 2.98 ms, repetition time (TR) = 2300 ms, inversion time (TI) = 900 ms, bandwidth = 240 Hz, acquisition matrix = 256 × 240 × 160, and voxel size = 1.0 × 1.0 × 1.0 mm. Neuromelanin-sensitive MRI was performed to assess contrast in the locus coeruleus. This sequence was aligned perpendicularly to the axis of the brainstem, guided by the whole-brain MPRAGE acquisition. Imaging parameters for the neuromelanin-sensitive sequence were flip angle = 20°, echo time = 1.5 ms, repetition time = 62 ms, and voxel size = 0.67 × 0.67 × 1.34 mm.

#### EEG Acquisition

Participants underwent a protocol involving EEG during a 5-minute eyes-open and 5-minutes eyes-closed resting-state period, followed by a short break and an auditory oddball task. The task design and behavioral data collection were implemented using MATLAB R2021b. High-density EEG data were recorded continuously using a 256-channel HydroCel Geodesic Sensor Net (Electrical Geodesics, Inc., EGI). Prior to recording, the cap was sized and soaked in a potassium chloride solution, following the manufacturer’s guidelines. EEG signals were acquired using EGI Net Station Acquisition 5.4.3, sampled at 1000 Hz, and amplified via the Net Amps 400 system. Electrode impedance was verified to be below 100 kΩ for all channels in accordance with EGI recommendations. Electrodes located on the cheeks and neck were excluded offline, leaving 191 cortical sensors for subsequent processing and analysis.

#### Resting-state EEG

Participants completed a resting-state EEG protocol comprising two consecutive conditions: a 5-minute eyes-open (EO) session followed immediately by a 5-minute eyes-closed (EC) session. During the EO condition, participants were instructed to maintain fixation on a centrally presented cross while keeping their eyes open. Throughout both sessions, they were asked to remain awake and alert while allowing their thoughts to wander freely. Only the eyes-closed recordings were used in the analyses reported in this study.

#### Task EEG Session

The oddball task consisted of auditory stimulus presentation, in which participants listened to sequences of spoken letters (B, K, and M) delivered through the monitor’s speakers. One of the three letters was designated as the standard stimulus, occurring in 60% of trials, while the remaining two served as oddball stimuli, each occurring in 20% of trials. Stimuli were presented with an inter-stimulus interval randomly jittered between 2.3 and 2.7 seconds. The presentation order was pseudorandomized, ensuring that no more than seven consecutive standard trials and no more than three consecutive oddball trials occurred sequentially.

The task was divided into three experimental sessions, with short breaks of up to one minute between sessions. Each session contained two blocks of 45 trials, separated by a 10-second inter-block break. The first two trials of each session were excluded from analysis, resulting in a total of 268 trials. Participants were instructed to maintain fixation on a centrally presented cross throughout the task. The first session was a passive listening condition in which participants were not required to respond to stimuli. In the second and third sessions, participants were instructed to respond via button press to a predefined target letter while ignoring the remaining non-target stimuli. In one of these two sessions, the frequently occurring standard stimulus served as the target (frequent session), whereas in the other, one of the oddball stimuli served as the target (oddball session). In the oddball session, the remaining oddball stimulus functioned as a non-target. The assignment of letters as targets, standards, or oddballs was counterbalanced across participants using a Latin Square design to minimize order effects and potential stimulus-specific biases. Spoken letter stimuli were used instead of pure tones, as high-frequency tones are often more difficult to perceive, particularly in older adults. The letters B, K, and M were selected based on their distinct auditory profiles, ensuring ease of differentiation. Auditory stimuli were generated using Audacity (an open-source digital audio editor), with sound volume and stimulus duration standardised at 300ms. Environmental conditions, including stimulus volume, screen brightness, and room illumination, were maintained consistently across all participants. Written instructions detailing the required response to stimuli were displayed at the beginning of each block, and verbal reminders were provided when necessary, with a maximum of two reminders per session if participants demonstrated consistent response errors.

### Derivation of Unit Phase Vector Field

#### Manifold approximation

To define locality on the manifold ℳ, we construct a proximity graph based on the point cloud *X*. For this purpose, we employ the continuous k-nearest neighbour (ck-NN) algorithm 110, which improves upon the classical k-NN approach by incorporating a local kernel density estimation. This method effectively adjusts for variations in sampling density across ℳ. In the ck-NN framework, two points i and j are connected if the condition 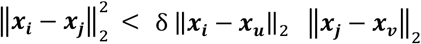 is satisfied. Here, u and v represent the k-th nearest neighbours of i and j, respectively, and ‖ ⋅ ‖_2_ denotes the Euclidean distance. The parameter δ is a scaling factor that governs the number of nearest neighbors, thereby influencing the extent of the local neighborhood.

#### Unit phase vector

Instantaneous phase ø(*x, y, z, t*) was extracted from band-pass filtered EEG signals (2–8 Hz, 9–20 Hz, 21–35 Hz) using the Hilbert transform. The complex phasor *ψ* = *e*^*i*φ^ was constructed, and spatial derivatives were estimated with manifold-based gradient filters (see *Gradient filters* in Methods). The phase gradient was recovered from the phasor derivative via 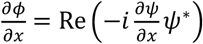, and analogously for y, z,

Where *ψ*\* is the complex conjugate of *ψ*. The result yields gradient components (∂φ/∂*x*, ∂φ/∂*y*, ∂φ/∂*z*). The norm of this gradient, |∇φ|, quantifies the local spatial rate of phase change. The temporal derivative |∂φ/∂*t*| was computed directly from the unwrapped phase time series. Combining spatial and temporal derivatives gives the local wave speed according to the eikonal relation:

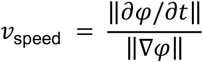

Multiplying this speed by the normalised negative gradient yields the phase velocity vector field:

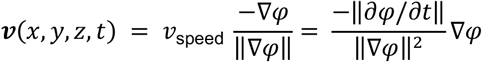

Here we consider only the direction of phase propagation, thus the vectors were further normalised to unit length:

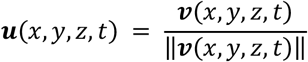

which we refer to as the unit phase vector field.

#### Gradient filters

Gradient filters are used to approximate the variation of scalar and vector fields on a manifold. These filters are applied twice in our analysis: first, to compute the phase vector vector field from scalar phase data using the 3D global coordinate frame, and second, to compute the directional derivatives of the vector field using local 2D coordinate frames defined on the manifold ℳ.

#### Gradient Filters for Scalar Fields

To construct gradient filters for the instantaneous phase data, our gradient filters operate in the global 3D coordinate frame. This allows us to compute the phase vector field directly in the ambient 3D space based on the local neighbourhood of each EEG channel. The directional derivative along the q-th axis (e.g., x, y, or z) is approximated using a weighted sum over the neighbours of a node

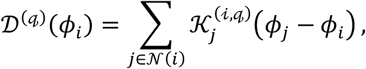

where 𝒩(*i*) is the set of neighbours of node i, 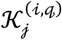 is the weight of the filter, dependent on the geometry of the graph and the spatial relationship between nodes i and j. By computing the directional derivatives along the three axes of the global 3D coordinate frame, the full gradient of the scalar field ø(*x, y, z, t*) is obtained as

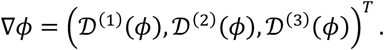

This computation provides a 3D vector field ∇ø at each node, representing the spatial rate of change of the scalar field.

#### Gradient Filters for Vector Fields

To compute directional derivatives of the 3D phase vector field *v* on the 2D manifold, we transition to local 2D coordinate frames at each point on the manifold. Each local frame *T*_*i*_ defines two orthogonal unit vectors 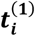 and 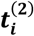 spanning the tangent plane (see tangent spaces in Methods). The vector field is projected onto the local frame:

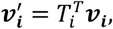

reducing the dimensionality from 3D to 2D. The variation of the vector field around node i is then approximated using a Taylor-series expansion in the local coordinates:

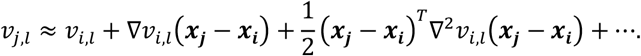

The gradient filters numerically approximate the first-order derivative operator ∇*v*_*i,l*_ as:

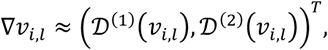

where 𝒟^(*q*)^Z*v*_*i,l*_[represents the directional derivative along the q-th unit vector in the local 2D frame. The directional derivative 𝒟^(*q*)^Z*v*_*i,l*_[is computed as:

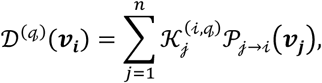

where 𝒦^(*i,q*)^ is the directional derivative filter acting along 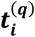, and *P*_*j*→*i*_ is the parallel transport operator. Parallel transport ensures that the computations are independent of the local curvature of the manifold.

#### Tangent spaces

We construct tangent spaces 𝒯_*i*_ℳ at each point i on the manifold. These tangent spaces linearly approximate ℳ within a local neighborhood. Specifically, the tangent space at a point 𝒯_*i*_ℳ is spanned by edge vectors *e*_*ij*_ ∈ *R*^d^, which point from i to its neighbouring nodes j on the proximity graph. To define the tangent space, we select *K* ≈ deg(*i*) neighbouring nodes, where K is a tuneable hyperparameter. The orthonormal basis for 𝒯_*i*_ℳ is derived from the m largest singular values 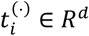 of the matrix formed by column-stacking the edge vectors *e*_*ij*_. This basis, *T*_*i*_ ∈ *R*^d×*m*^, spans the tangent space:

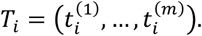

Projecting a signal *f*_*i*_ to the tangent space is then accomplished by 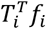, which minimizes the projection error in the I_2_ sense. These computations are performed using a modified Parallel Transport Unfolding package^111^ and the MARBLE package freely available on github (github.com/agosztolai/MARBLE)^112^

#### Connections

Once local tangent spaces and their orthonormal bases are established, we define a parallel transport map *P*_*j*→*i*_ to align the local frame at node j with the frame at node i. This alignment is essential for computing the directional derivatives in a common space. Although parallel transport is generally path-dependent, for neighboring nodes i and j that are sufficiently close, we assume the unique minimal rotation - the Levy-Civita connection. The parallel transport operator *P*_*j*→*i*_ is represented as the orthogonal transformation matrix *O*_*ji*_, which satisfies

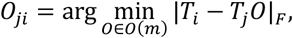

where | ⋅ |_*F*_ denotes the Frobenius norm. The matrix *O*_*ji*_ is computed using the Kabsch algorithm 113 which provides a unique solution (up to a change of orientation) to this optimisation problem.

#### Curl and divergence

To analyse the rotational and compressional characteristics of the vector field *v* on the 2D manifold, we compute its curl and divergence. These quantities are derived using the Jacobian matrix of the phase vector field, which encodes its local variation. The Jacobian matrix *J*_*i*_ at each point i on the manifold is defined as

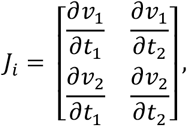

where (*t*_1_, *t*_2_) are the two orthogonal directions of the local tangent space defined by the 2D local coordinate frame *T*_*i*_, *v*_1_ and *v*_2_ are the components of the velocity field *v*_*i*_ in the local coordinate frame. The entries of *J*_*i*_ are computed using the directional derivative filters 𝒟^(*q*)^, as described in Gradient Filters.

The divergence of the velocity field at node i is computed as the trace of the Jacobian matrix:

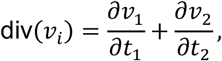

measuring the net rate of expansion or compression of the phase vector field at the point i. The curl in two dimensions is the antisymmetric part of the Jacobian:

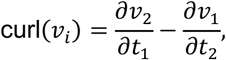

measuring the rotational tendency of the phase vector field around the point i. Importantly, since the curl and divergence are scalar values, they are independent of the local tangent plane orientation. The connections between tangent planes provides a locally linear space in which to determine the directional derivatives and hence the Jacobian (or higher-order derivatives such as the Hessian).

### Derivation of Routing Modes

To derive the fundamental modes of divergence and vorticity in the EEG signal, we first constructed a data matrix for each participant by concatenating all time-resolved divergence or vorticity maps (time snapshots) into a single matrix. For each subject, this matrix contained one row per time point and one column per EEG sensor, capturing the full spatiotemporal evolution of the wavefield. PCA was then applied separately to each subject’s matrix to extract a set of orthogonal spatial modes, which we refer to as subject-level eigenmodes. These modes were obtained by performing eigenvalue decomposition on the covariance matrix of the concatenated divergence or vorticity maps. The resulting eigenvectors represent a reduced basis of spatial patterns—ordered by the amount of variance they explain—that characterise dominant modes of cortical flow in each individual. To derive group-level modes, we constructed a second matrix by vertically concatenating the time-resolved divergence or vorticity maps across all participants. PCA was then applied to this pooled matrix to extract a shared set of group-level eigenmodes that capture the principal spatial patterns common across subjects. To compute how these modes evolve over time— whether at the subject or group level—we projected the time-resolved EEG divergence or vorticity maps onto the corresponding subject or group level eigenmodes. This projection was performed by computing the dot product between the eigenmodes and the EEG-derived wavefield at each time point, yielding a time series that quantifies the instantaneous activation strength of each mode.

### Kuhn–Munkres algorithm

To optimise the alignment of routing modes between two sets—such as group-level and subject-specific routing modes—we employed the Hungarian matching algorithm, also known as the Kuhn–Munkres algorithm. This method establishes a one-to-one correspondence between routing modes from the two sets that maximises the total absolute correlation across all matched pairs. Each set of routing modes was represented as a matrix of size N × N, where rows corresponded to the weight at an individual EEG sensor and columns to an individual routing mode. Matrix ***A*** contained, for example, the group-level routing modes, and matrix ***B*** the subject-specific routing modes. The objective was to permute the columns of ***B*** such that the alignment with ***A*** maximised the total absolute correlation between corresponding routing modes. To solve this assignment problem, we first computed the pairwise absolute correlations between all columns in matrices ***A*** and ***B***, resulting in an *N* × *N* correlation matrix. We then constructed a cost matrix by taking the negative of these absolute correlations. Minimising the total cost is mathematically equivalent to maximising the total absolute correlation. The Hungarian algorithm was then applied to identify the optimal permutation of columns in matrix ***B*** that maximised alignment with matrix ***A***. Once the optimal matching was identified, the columns of the subject-specific eigenmode matrix were reordered according to the derived permutation. This procedure enabled consistent comparison of routing mode structure and dynamics across individuals.

### Null Model for Mode-to-Mode Correlation

For each comparison between two routing mode families (e.g., morning vs. evening routing modes), we generated subject-level surrogates to estimate the mode-to-mode correlations expected by chance. For a given subject’s divergence and vorticity maps we preserved the electrode montage but replaced each channel’s time series with that from a randomly selected donor participant. This procedure preserved marginal per-channel statistics while disrupting cross-channel temporal dependencies that generate coherent flow structure. Eigenmode decomposition was then repeated on the surrogate data, and the absolute Pearson correlation between each surrogate mode and the corresponding reference-family mode was computed. This resampling and decomposition were repeated 1,000 times, yielding a null distribution of 1,000 correlation values for each routing mode. The emperical correlation coefficient was deemed significant if it exceeded the 95th percentile of its mode-specific null distribution (*P* < 0.05), with Bonferroni correction applied across the 50 tested routing modes (corrected threshold = 0.05/50).

### Cross-mode covariance (CMC)

For each participant, time-resolved divergence and vorticity fields were projected onto the subject-specific eigenmodes derived from the decomposition, yielding a mode-specific amplitude matrix *A* (rows = time points, columns = modes). Each column of *A* was z-scored (zero mean, unit variance), and the cross-mode covariance matrix (CMC) was computed as *CMC* = *A*^*T*^*A*. To quantify similarity between participants or sessions, we extracted the lower triangular elements (excluding the diagonal) and unwrapped them into a one-dimensional vector. Empirical similarity was then evaluated by computing the cosine similarity between unwrapped CMC vectors (e.g., subject 1 vs. subject 2; morning vs. evening). Statistical significance was assessed by generating a surrogate null distribution: CMC vectors were randomly permuted, cosine similarity was recomputed, and the procedure was repeated 5,000 times. Observed cosine similarity values were z-scored relative to the null distribution (mean and standard deviation), and group-level effects were tested with two-tailed one-sample t-tests of the z-scores against zero, with positive values indicating similarity greater than chance.

### Routing mode activation on single trials

To estimate mode activations on a trial-by-trial basis, we first computed time-resolved divergence and vorticity maps for each trial during the frequent, oddball, or passive listening sessions. These spatiotemporal maps were then projected onto each participant’s task-specific divergence or vorticity eigenmodes, yielding a time series of mode activations for each trial (90 trials per session). This projection captured the dynamic expression of each routing mode from stimulus onset to one second post-stimulus. Because routing mode activations typically oscillated between positive and negative amplitudes, we quantified activation strength by computing the variance of each routing mode’s amplitude time series over the one-second post-stimulus window. This procedure yielded a trial-by-routing mode feature matrix of size 90 × 25 per task session (Passive, Frequent and Oddball sessions), where each row represented a single trial and each column corresponded to the amplitude variance of one of 25 routing modes.

### Classification of task session using random forests

To test whether distinct cognitive functions rely on different routing architectures, we performed single-trial classification between pairs of task sessions—Passive listening (P) versus Frequent (F), P versus Oddball (O), and F versus O. Classifiers were implemented in MATLAB using the *TreeBagger* function with 200 decision trees, each constrained to a maximum depth of three splits. Model evaluation used five-fold cross-validation with an 80:20 train– test split, and performance was quantified as the area under the receiver operating characteristic curve (AUC) on the held-out test data. Feature importance was estimated using MATLAB’s *oobPermutedPredictorImportance* function. This method quantifies how strongly each predictor contributes to classification by measuring the increase in out-of-bag (OOB) error when its values are randomly permuted across the OOB samples for each tree. For each predictor, the importance score is defined as the mean increase in error (relative to the baseline OOB error), normalised by its standard deviation across the ensemble. Higher scores therefore indicate predictors whose perturbation most strongly degrades model performance, providing a relative measure of which routing eigenmodes contributed most to task discrimination. Separate classifiers were trained on low-, mid-, and high-frequency band activations, yielding six binary classifiers in total (three for divergence and three for vorticity eigenmodes).

### Partial least squares analysis

We applied partial least squares (PLS) regression to identify multivariate associations between mode rerouting rates and regional grey-matter volume. Analyses were conducted in MATLAB using the myPLS toolbox (https://github.com/valkebets/myPLS-1). For each participant, rerouting rate was defined as the zero-crossing frequency (sign-changes per second) of the amplitude time series for the first 25 low-frequency (LF) divergence and vorticity modes during rest. The predictor matrix *X* therefore comprised rerouting rates for 50 modes per subject (dimensions: 30 subjects × 50 routing modes). The response matrix *Y* contained regional grey-matter volumes from the Schaefer 200-parcel atlas, extracted from T1-weighted MRI scans using FreeSurfer’s recon-all. Subcortical volumes from 14 bilateral structures (caudate, hippocampus, putamen, thalamus, pallidum, nucleus accumbens, amygdala) were included, and all regional values were normalised to total intracranial volume. Both *X* and *Y* were z-scored across participants to remove scale differences.

PLS identifies latent variables (LVs) that maximise covariance between XX and YY by singular value decomposition of the cross-covariance matrix:

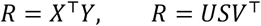

where UU and VV are left and right singular vectors (saliences for XX and YY), and SS contains the singular values. Subject-specific PLS scores were obtained by projecting the data onto the salience vectors:

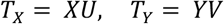

Permutation testing (5,000 iterations) established statistical significance by recomputing singular values after permuting the rows of *Y*. The empirical *P*-value for each LV was defined as the proportion of permutations exceeding the observed singular value. Stability of saliences and loadings was assessed by bootstrap resampling (500 iterations), with Procrustes rotation to align resampled solutions to the original. Confidence intervals and standard errors were derived from the bootstrap distributions.

## Funding

UK Dementia Research Institute

Engineering and Physical Sciences Research Council (EPSRC), award number (EP/L016737/1).

We are grateful to M. Desforges and Y. Gal-Shohet for the stimulating and thoughtful discussions.

We also wish to thank S.L. Brunton and B.W. Brunton, as well as J.Roberts and the co-authors of *Metastable Brain Waves*, who inspired this project.

## Author information

These authors contributed equally: Matteo Vinao-Carl, Robert L. Peach.

## Contributions

M.V.C. and N.G. conceived the study and formulated the overarching research goals. M.V.C. developed the methodology, performed validation, conducted the formal analyses, and wrote the original draft. M.V.C., R.L.P. and A.G. implemented and tested the software. V.J. and I.R.V. (cohort 1), and M.C.B.D., E-J.M. and D.J.S. (cohorts 2 and 3) performed the investigations and provided patient resources. D.K. curated the structural MRI metadata. M.V.C. prepared the visualizations. M.V.C., R.L.P. and N.G. critically reviewed and edited the manuscript. Supervision was provided by R.L.P. & N.G., with project administration by N.G. Funding was acquired by M.V.C. and N.G.

## Corresponding Author

Correspondence to Matteo Vinao-Carl or Nir Grossman.

## Data and materials availability

The raw data for cohort 1 can be obtained from the following repository at zenodo; zenodo.org/records/10663994. The raw data for cohorts 2 and 3 (healthy older adult and Alzhiemer’s disease cohorts) were collected by the authors at the UK DRI CR&T and can be requested by contacting D.J.S., under sufficient material transfer agreements. The analysis scripts, algorithms, as well as the final data and code for generating the figures, will be made available upon publication.

