## Supplementary Materials for "A universal dynamic routing architecture governs the flow of human cortical activity"

#### Supplementary Figures

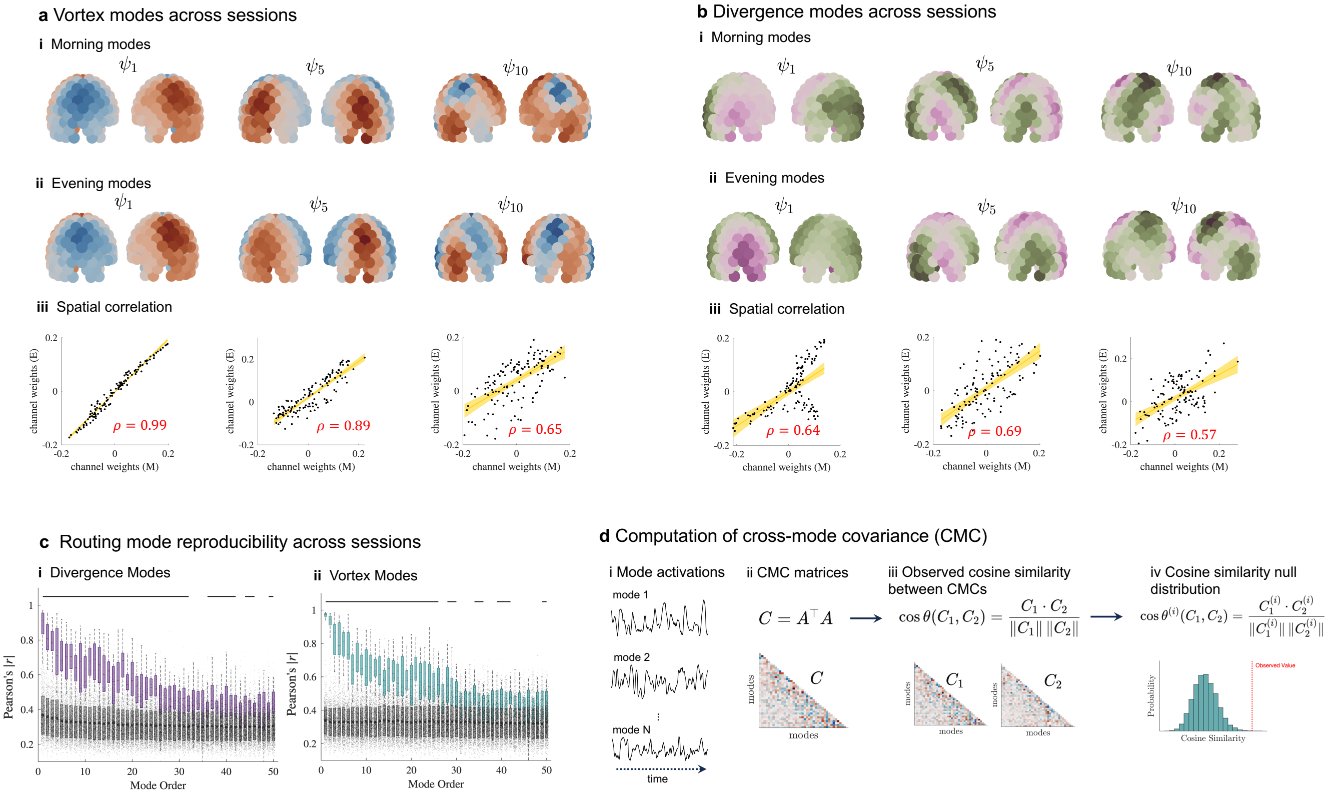


Fig.S1. (related to Fig.2).

*The Healthy Young Cohort is from the overnight sleep study reported in Jaramillo et al., 2024 [1].*

**a,** α-band vorticity routing eigenmodes from a representative participant recorded in the morning (i) and evening (ii), together with channel-wise spatial correlations between eigenmodes $\psi_1$, $\psi_5$ and $\psi_{10}$ across morning and evening sessions (iii).

**b,** Same as **a** for divergence modes.

**c,** Within-subject spatial reproducibility of eigenmodes. Box plots show the distribution of absolute Pearson correlation coefficients between morning and evening eigenmodes for modes $\psi_{1\text{-}50}$ across all participants. Boxes denote the inter-quartile range (25th–75th percentiles); central line, median; whiskers, the most extreme values excluding outliers. Grey shaded boxes depict the null distribution in which local spatiotemporal statistics are preserved but long-range structure is disrupted. Solid black bars indicate modes that remain significant after Bonferroni correction for 50 comparisons ($P < 0.05$).

**d**, Cross-mode covariance (CMC) pipeline. i, Mode-specific activation time series were obtained by projecting time-resolved divergence or vorticity fields onto their corresponding group-level eigenmodes. ii, For each participant, we z-scored each mode's amplitude time series for the first 50 modes to form matrix $A$ (time points × normalised eigenmodes). iii We computed the cross-mode covariance (CMC) matrix $C = A^T A$. iii, Similarity between CMCs from different sessions from the same participant ($C_{morning,}$ $C_{evening}$) was quantified with cosine similarity. iv Significance was assessed by z-scoring the observed cosine similarity against a surrogate null distribution obtained by independently permuting the columns of $C_{morning}$ and $C_{evening}$ and recomputing the similarity 5,000 times.

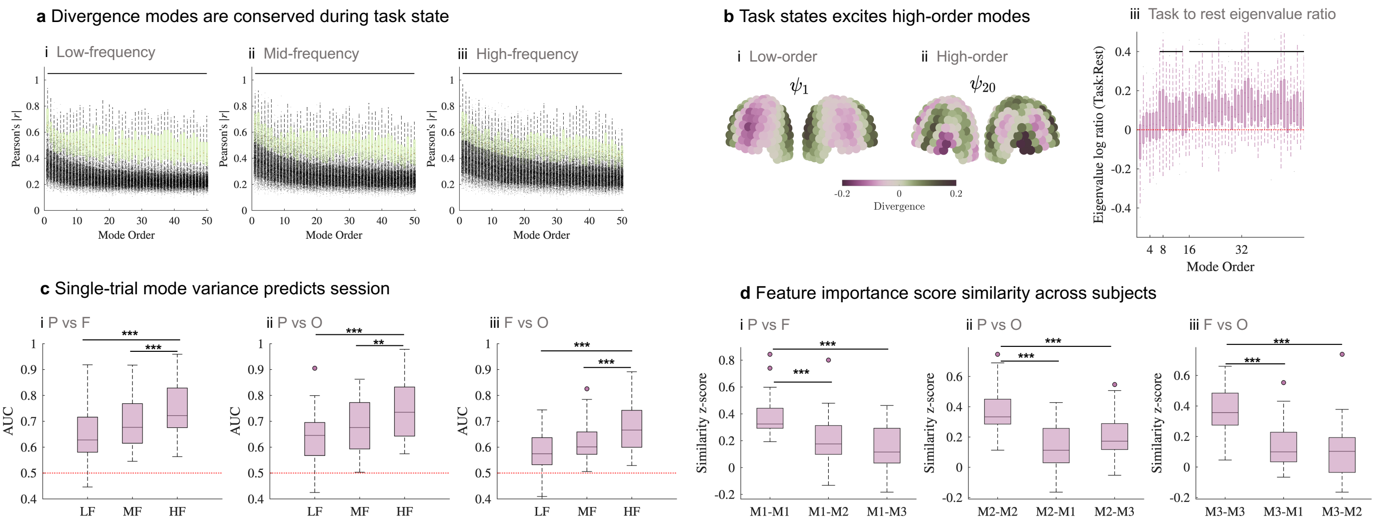


### Fig.S2. (related to Fig.4)

**a Conservation of divergence eigenmodes during task state.** Absolute Pearson correlations between task-state divergence modes and the resting-state LF family (reference family) after re-ordering using the Hungarian algorithm. Green boxplots show the inter-quartile range for low- (i), mid- (ii) and high-frequency (iii) bands; whiskers denote most extreme values. Black boxplots depict null-model distributions, and solid black bars mark modes whose correlations exceed the surrogate (Bonferroni-corrected for 50 comparisons, $P < 0.05$).

**b High-order mode amplification at high frequencies.** i Group-level low-order divergence mode ($\psi 1$). ii Group-level high order divergence mode ($\psi 20$). iii $\text{Log}_{10}$ ratios of divergence-mode eigenvalues in task versus rest for the first 50 eigenmodes, derived from HF activity. Positive values indicate excitation of a given mode during the task relative to rest, negative values indicate suppression. Significant departures from zero (paired two-tailed *t*-test, $P < 0.05$ uncorrected; black bars) appear for higher-order modes ($\psi > 12$), indicating preferential routing of HF activity through these modes during the task.

**c Single-trial decoding.** Binary-classification accuracy (mean ± 95 % CI) based on mode-amplitude variance for 90 trials per session: passive vs frequent (i), passive vs oddball (ii) and frequent vs oddball (iii). Classification for all contrasts exceed chance, with HF features yielding the highest performance.

**d Shared feature profiles across subjects.** Subjects exhibited similar feature-importance vectors from the random-forest models. For each contrast—PvF (i), PvO (ii) and FvO (iii)—similarity is higher for matched models (same behavioural rule, different subjects) than for mismatched models (different behavioural rule, different subjects), indicating that identical task instructions recruit comparable subsets of divergence modes across individuals.

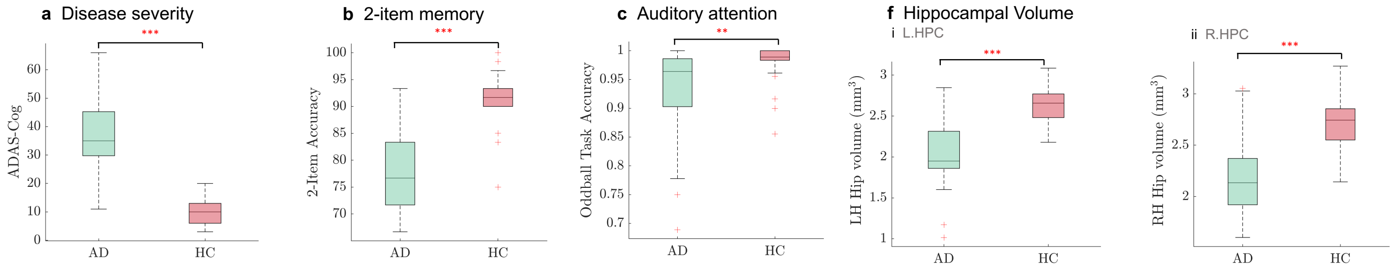


#### Fig.S3 (related to Fig.5).

**a**, Alzheimer's Disease patients (AD, green) exhibit significantly higher scores on the Alzheimer's Disease Assessment Scale–Cognitive Subscale (ADAS-Cog[2]) compared to healthy controls (HC, red); unpaired two-tailed *t*-test, $t = 11.12$, $P = 4.17 \times 10^{-16}$).

**b**, Performance on a 2-item memory task is significantly impaired in AD patients relative to HC ($t = 6.57$, $P = 3.4 \times 10^{-8}$).

**c**, AD patients show reduced performance on the auditory attention task, indexed by mean hit rate averaged across frequent and oddball sessions ($t = -2.90$, $P = 0.005$).

**d**, Hippocampal volume is reduced in AD relative to HC in both the left (**i**, $t = -6.77$, $P = 1.17 \times 10^{-8}$) and right hippocampus (**ii**, $t = -5.66$, $P = 6.59 \times 10^{-7}$).

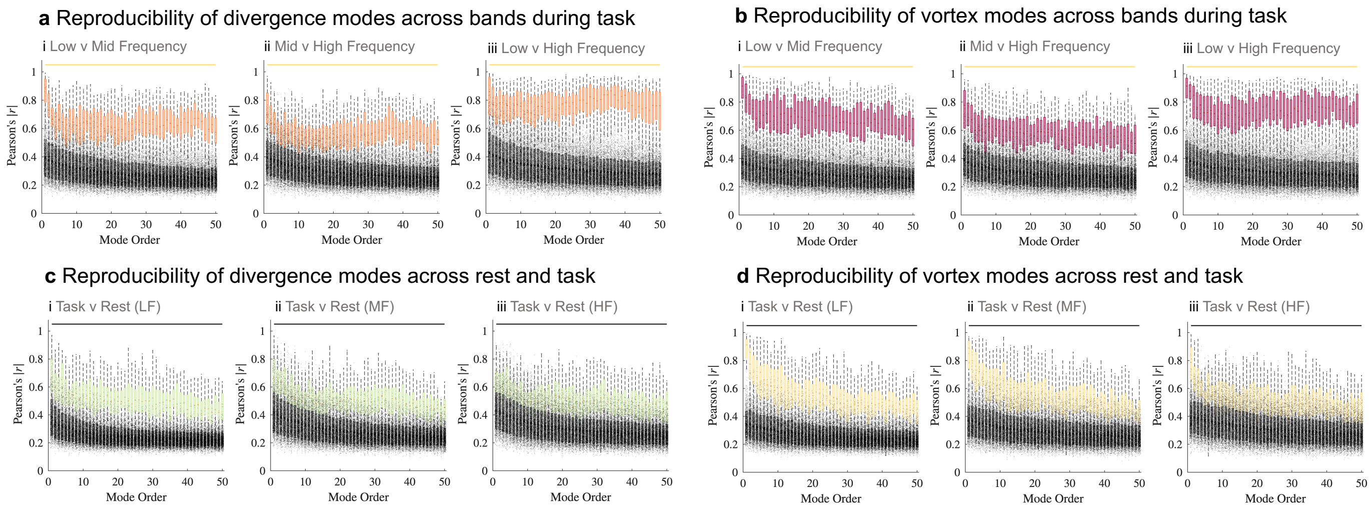


### Fig.S4. (related to Fig.5)

**a–b Across-band reproducibility.** Absolute Pearson correlation coefficients between subject-level modes computed from different frequency bands within a subject: LF vs MF (i), LF vs HF (ii), and MF vs HF (iii). Cross-band similarity is shown separately for divergence (**a**) and vortex (**b**) modes. All modes were reordered relative to the reference family (LF resting state group-level modes).

**c–d Cross-state reproducibility.** Similarity of task modes to resting-state LF modes (Reference Family) for divergence (**c**) and vortex (**d**) modes. Comparisons are shown for LF vs Reference Family (i), MF vs Reference Family (ii), and HF vs Reference Family (iii). Boxplots indicate the interquartile range; whiskers extend to the most extreme non-outlier values. Yellow lines denote modes with significantly greater similarity than expected under the null distribution (black boxplots), Bonferroni-corrected for 50 multiple comparisons ($P < 0.05$).

2. Kueper, J. K., Speechley, M. & Montero-Odasso, M. The Alzheimer's Disease Assessment Scale–Cognitive Subscale (ADAS-Cog): Modifications and Responsiveness in Pre-Dementia Populations. A Narrative Review. J Alzheimers Dis 63, 423–444.
